# High Dimensionality Reduction by Matrix Factorization for Systems Pharmacology

**DOI:** 10.1101/2021.05.30.446301

**Authors:** Adel Mehrpooya, Farid Saberi-Movahed, Najmeh Azizizadeh, Mohammad Rezaei-Ravari, Farshad Saberi-Movahed, Mahdi Eftekhari, Iman Tavassoly

## Abstract

The extraction of predictive features from the complex high-dimensional multi-omic data is necessary for decoding and overcoming the therapeutic responses in systems pharmacology. Developing computational methods to reduce high-dimensional space of features in *in vitro, in vivo* and clinical data is essential to discover the evolution and mechanisms of the drug responses and drug resistance. In this paper, we have utilized the Matrix Factorization (MF) as a modality for high dimensionality reduction in systems pharmacology. In this respect, we have proposed three novel feature selection methods using the mathematical conception of a basis for features. We have applied these techniques as well as three other matrix factorization methods to analyze eight different gene expression datasets to investigate and compare their performance for feature selection. Our results show that these methods are capable of reducing the feature spaces and find predictive features in terms of phenotype determination. The three proposed techniques outperform the other methods used and can extract a 2-gene signature predictive of a Tyrosine Kinase Inhibitor (TKI) treatment response in the Cancer Cell Line Encyclopedia (CCLE).

**Key Points:**

- Matrix Factorization (MF) is a useful framework for high dimensionality reduction in systems pharmacology.
- Novel feature selection methods using the incorporation of the mathematical conception of a basis for features into MF increases the performance of feature selection process.
- Feature selection based on the basis-concept in MF can provide predictive gene signatures for therapeutic responses in systems pharmacology.

## 1 Introduction

One of the challenges in the era of systems medicine is to decode and select personalized features for an individual patient to design personalized therapeutic regimens [1, 2], even if the disease has been modeled analytically [3, 4, 5, 6]. Although systems pharmacology-based mechanistic models which are built based on the known interactions among molecular mechanisms and phenotypes provide a path to solve this problem, but high dimensionality of clinical and molecular characteristics, including multi-omic features, adds more complexity to uncovering the mechanisms causing complex diseases in any model system and patient [7]. Systems-level study of interaction networks constructed from omics data can lead to static models connecting molecular data to phenotypes [8]. Yet, these types of models are still high-dimensional and lack temporal characteristics of disease evolution. A path to design personalized molecular sets assigned to a specific phenotype of a complex disease like cancer is to use phenomenological approaches to reduce the high-dimensional data [9]. The emergence of molecular signatures on this domain requires new methodologies for both model reduction and feature selection in large molecular omic and multi-omic data sets. In systems pharmacology, predicting the therapeutic responses in *in vitro* and *in vivo* models and detecting the molecular signatures related to drug resistance help to narrow down the therapeutic regimens for testing in clinical trials [1, 10]. Highdimensional data has noise and redundancy and includes irrelevant information controlled by some unknown parameters required for the model to run. For this reason, analysis of large biological data sets does not always end in robust models and is computationally costly while being prone to over-fitting [11, 12]. In addition, if the modeling process is selected to stand on the distance between molecular features, the designed methods face even more performance degeneration due to adverse impacts of the high-dimensional feature space on the calculation of feature distances. With the emergence of advanced information technology and its increasing applications in electronic communications, a wide variety of modern devices have been designed based on big data whose dimension is exceptionally high [13, 14, 15]. The noise and redundancy, including the irrelevant information, come from a large number of complex parameters in the model, and it results in a limited ability to generalize the learning model and impose inefficient requirements such as more memory usage and longer processing time on the computational devices since the corresponding algorithms suffer enormously from over-fitting [16, 17, 18, 19, 20, 21].

To face the challenge of high dimensionality, two well-known approaches have been proposed. One of these methods is “Feature Extraction” which reduces the dimension by projecting the data given in a higher-dimensional space, through a proper mapping, into a subspace of this space whose dimension is lower enough than that of the original space [22, 23, 24, 25, 26]. The other technique is “Feature Selection” which reduces the high dimensionality by selecting a smaller number of features compared to the original feature set so that the selected features have the capability to represent the original feature space optimally [27, 28, 29]. It is proved that both techniques, particularly feature selection, substantially contribute to eliminate redundancy and noise, remove the irrelevant parameters and enhance the efficiency of data analysis computations, thus leading to a decrease in the costs associated with time and storage [30, 31]. This way, feature extraction, and feature selection provide considerable improvements in the clustering and classification accuracy, respectively. It should be pointed out that feature selection maintains the original features’ conceptual information by preserving the original features representation more perfectly compared to feature extraction, which constructs new features. For this reason, it is more common to use feature selection than feature extraction to develop novel algorithms for handling dimension reduction tasks in recent years [22, 32].

A broad and significant category of the applications of machine learning techniques discussed can be merged with systems pharmacology methodologies to uncover the quantitative features involved in the evolution and treatment of complex diseases such as cancers [1]. One of the first steps to build systems pharmacology models and platforms is to analyze big datasets such as gene expressions and transcriptomics. Since these datasets, considering phenotypes associated with them, are big and complex datasets, reducing their high dimensionality is a key to be able to use them in systems pharmacology [1]. For instance, to determine the sensitivity and resistance to a specific therapeutic *in vitro* and *in vivo*, it is necessary to explore the bio-molecular datasets such as gene expression profiles of hundreds of cancer cell lines [2]. Exploring these datasets to extract features related to the mechanisms causing therapeutic-resistant phenotype requires reducing the high-dimensional space of the molecular features. The complexity of the omic datasets, including gene expressions, originates from two major properties of the biological systems. The first one is intrinsic geometry. It indicates the geometrical properties of these data need to be preserved when the features initially exist in a manifold of high-dimension mapped into a lower-dimension sub-manifold of the base space. The other characteristic of such big data is the noise and enormous amount of outliers.

Over the past decades, subspace learning techniques have demonstrated a considerable ability to deal with feature selection problems and their applications to the analysis of complex biological systems [12, 33, 34, 35, 36]. It is required that the subspace learning techniques and feature selection methods, particularly when they are employed to handle gene expression data, are equipped with some proper and efficient tools. In this regard, some modifications have been incorporated into these methods. One good example is subspace learning for unsupervised feature selection via matrix factorization (MFFS), introduced in [37], in which a subspace learning framework is designed from the perspective of the matrix decomposition. The MFFS framework and the idea of minimum redundancy were merged to adapt unsupervised feature selection via maximum projection and minimum redundancy (MPMR) [38]. Another interesting example is subspace learning-based graph regularized feature selection (SGFS), investigated in [39], which can maintain the features intrinsic information in terms of features geometry. Then utilizing matrix factorization techniques again, an unsupervised method, feature selection based on maximum projection and minimum redundancy (MPMR), was explored. All these algorithms can provide a highly accurate estimate of the feature space by learning a proper subspace and also produce satisfactory outcomes for the corresponding feature selection task.

Although the mentioned methods provide powerful tools to successfully develop new practical models for dimension reduction problems, they have not been vastly applied to systems pharmacology. The major goal of this paper is to establish a number of novel feature selection frameworks and apply them to cancer gene expression data. In the proposed methods, not only are the notable and outstanding achievements of the MFFS, MPMR and SGFS methods considered, but also the performance of the introduced methods is improved in terms of classification and clustering. For this, the notion of a basis, which is an essential concept in the theory of linear vector spaces, is applied. Normally, a basis possesses two fundamental characteristics, each of which has a positive and constructive impact on the discriminative feature selection process [35]. The first property is that a basis generates the whole feature space making use of the basis elements. This means that the characteristics of original features are included in the basis whose number of elements is far smaller than that of the whole feature space. A major advantage in this case is that rather than considering the original features, a basis can be utilized to represent them. It leads to a considerable reduction in the number of the selected features and the calculating processes. The second property of a basis is its elements that are linearly independent, which indicates a significant decrease in the data redundancy.

Figure 1 illustrates a schematic diagram of the major set of concepts used in the proposed methods. This figure shows that considering an original feature data, a basis of features is primarily built for the original feature space. Accordingly, the feature selection problem is defined based on the constructed basis. Through some regularizers such as “the *L*_2,1_-norm sparsity regularization”, “the redundancy of the basis space” and “the manifold of the basis space”, three feature selection methods are introduced under these regularizations, and the feature matrix **G** is learned to enhance the feature learning performance.

**Figure 1:**
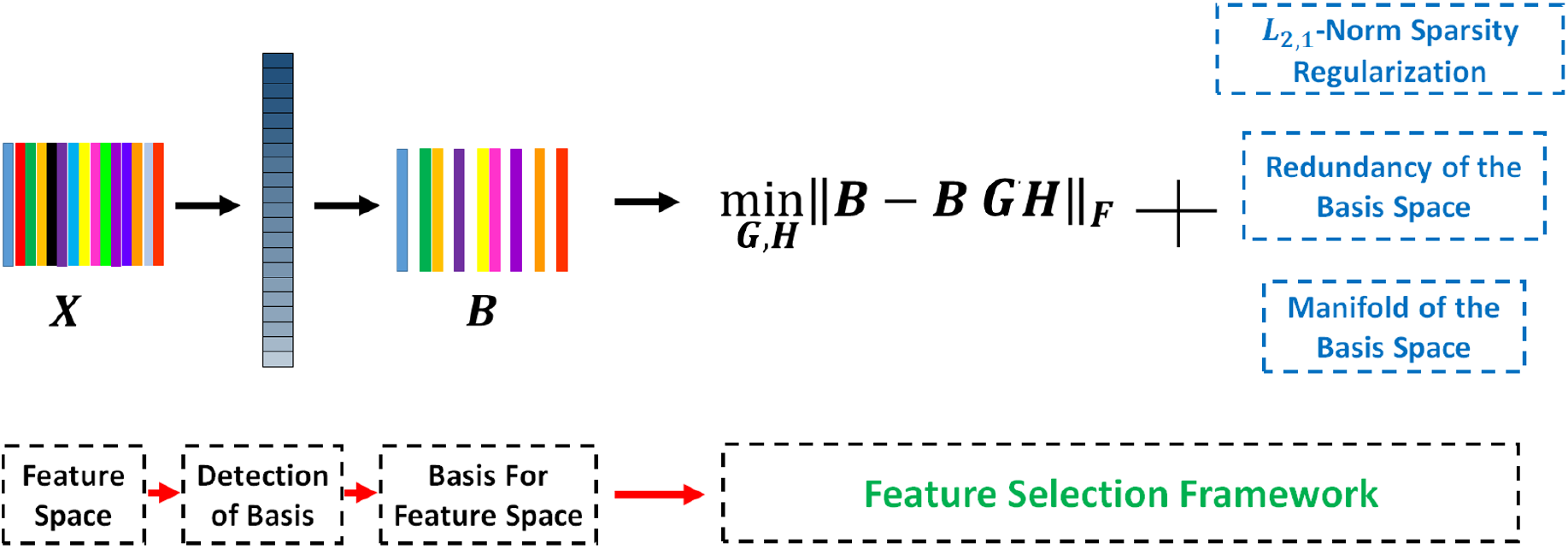
A schematic diagram of the fundamental framework of the proposed methods.

## 2 Motivation and Background

This section discusses the substantial role of a basis of features for the feature selection problems and describes an algorithm to construct a basis for the features space. Throughout the paper, the lowercase letters in Italic are used to denote the scalers, the uppercase letters in boldface represent matrices, the lowercase letters in boldface indicate vectors, and the uppercase letters in italic are used for sets. For a matrix **A**, the set including all linear combinations of its columns is denoted by span(**A**). The notation **E**_*k*_ represents the *k* × *k* identity matrix. For a given matrix **A, A**^*T*^ and Tr(**A**) denote the transpose and the trace operators, respectively. In addition, ∥**A**∥_*F*_ is the Frobenius norm of the matrix **A**. For a matrix **A** ∈ ℝ^*m*×*n*^, the *L*_2,1_–norm of **A** is given by 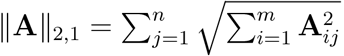.

The data matrix is represented by **X** = [**x**_1_; **x**_2_; … ; **x**_*n*_] = [**f**_1_, **f**_2_, …, **f**_*d*_] ℝ^*n*×*d*^, where *n* and *d* indicate the number of samples and that of features, respectively. It is also supposed that *n* ≤ *d*. For a feature subset *X*_*I*_, the corresponding submatrix of **X** is simply represented by **X**_*I*_ that is a matrix of order *n* × *k* in which *I* and |*I*| show the index set of the selected features and the number of indices of *I*, respectively. Given a vector space *V* of dimension *n*, a basis for *V* is a subset *B* = {**v**_1_, **v**_2_, …, **v**_*n*_} of *n* linearly independent elements of *V* such that *V* is spanned by the vectors of *B*. Given the matrix of features **X** = [**f**_1_, **f**_2_, …, **f**_*d*_], it is supposed throughout this article that the set *B* is composed of *n* elements and is a basis of the vector space spanned by **X** as well as assuming that rank(**X**) = *n*. The rank of a matrix **X**, rank(**X**), is the dimension of the vector space produced by the rows or the columns of **X**. It can also be proven that rank(**X**) is equal to the maximum number of rows or columns of **X** that are linearly independent.

### 2.1 Motivation

The notion of directional distance between two subspaces has been discussed in learning problems regarding subspaces and several feature selection methods have been established based on it.

#### Definition 1

(Directional Distance [37]). Given two matrices **S**_1_ ∈ ℝ^*m*×*n*^ and **S**_2_ ∈ ℝ^*m*×*p*^, the distance from 𝒮_1_ = span(**S**_1_) to 𝒮_2_ = span(**S**_2_) is calculated by:

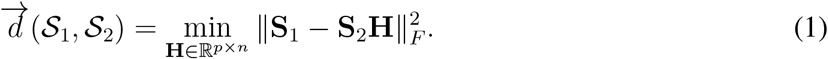

One most well-known and practical technique for feature selection is presented in Problem 1 that is rested on the directional distance defined above and has been the center of attention as the foundation of a number of powerful state-of-the-art algorithms in recent years [34, 35, 36, 37, 38, 39].

#### Problem 1.

Let **X** ℝ^*n*×*d*^ and *k* ∈ Z^+^. Considering ℐ as a feature index set for which |ℐ| = *k*, a feature selection problem is formulated by Eq. (2).

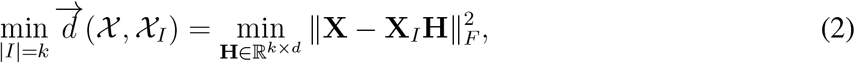

where 𝒳 = span(**X**) and 𝒳_*I*_ = span(**X**_*I*_).

Saberi-Movahed et al. [35] studied the importance of the representation of features provided by the notion of a basis of span(**X**) in terms of the improvements that this representation makes on the feature selection performance in Problem 1. Specifically, two highly significant properties were investigated as follows.

1. Let *B* be a basis of span({**f**_1_, **f**_2_, …, **f**_*d*_}) and B = span(**B**). Then

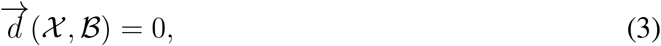
2. Let *X*_*I*_ ⊆ *X* be a feature subset. Then

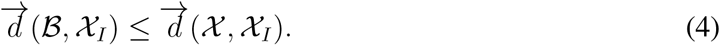

Item (1) states that for a basis *B* of a feature subspace 𝒳, its span, B, can be selected as a representation of 𝒳 since ℬ coincides with 𝒳. Furthermore, Item (2) indicates that the representation of 𝒳 with ℬ, i.e., assuming that =, encourages us to replace 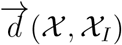 in Problem 1 by 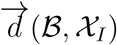 and it guarantees that this replacement will not make the results of data fitting worse than that of the original problem [35].

Considering the merits of a basis in feature selection problems, Saberi-Movahed et al. [35] proposed a new objective function, given by Eq. (5), in which the original feature space was represented by a basis of this space. Problem 2 summarizes this method which improved the method presented by Problem 1.

#### Problem 2.

Let *X* be the original set of features and *B* be a basis of the span of *X*. In addition,

ℬ = span(**B**) and 𝒳_*I*_ = span(**X**_*I*_). Assuming that ℐ is a set composed of *k* indices of the features, an objective function can be formulated as:

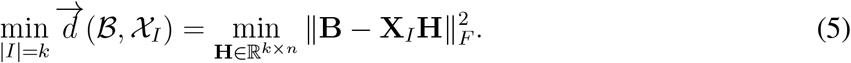

Recently, a supervised feature selection technique, SFS-BMF, rested on Problem 2, was presented in [35] so that a basis of features was applied to the selection process. The advantage of this method over a set of benchmark techniques was demonstrated by conducting diverse experiments on a number of gene expression datasets. Inspired by satisfactory outcomes from supervised feature selection founded on the notion of a basis, we investigate the impacts of this approach on the outcomes of feature selection problems in the unsupervised mode.

The main goal of the feature selection is to choose a feature subset to decrease the noise and to enhance the efficiency and efficacy of a learning task, such as classification, prediction and clustering [40, 41]. To diminish the dimension of feature space, the amount of the lost information needs to be as low as reasonable so that the obtained space can supply sufficient information for the learning task. In fact, it is essential to pick out features that are relevant for a learning task, but at the same time it is important to have a set of features which is not redundant for the purpose of enhancing the robustness [28]. In this regard, one of the most important issues in the process of feature selection is the redundancy. Redundancy evaluates how similar the features are to each other and how much inserting a new feature into a specific feature set plays an effective role in the data analysis [42].

In the context of linear algebra, there is a direct relationship between the redundancy and the notion of linear independence [43]. Specifically, a set of vectors {**x**_1_, **x**_2_, …, **x**_*d*_}in ℝ^*n*^ is called linearly dependent if there exist non-zero scalars *α*_1_, *α*_2_, …, *α*_*d*_ in ℝ such that *α*_1_**x**_1_ + *α*_2_**x**_2_ +…+ *α*_*d*_**x**_*d*_ = 0. This means that one of these vectors is a linear combination of some of the others. On the other hand, it is said that the vectors are linearly independent if they are not linearly dependent, that is, the expression *α*_1_**x**_1_ + *α*_2_**x**_2_ +…+ *α*_*d*_**x**_*d*_ = 0 implies that *α*_*i*_ = 0 for *i* = 1, …, *d*. For an example, let us consider the vectors

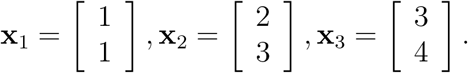

It can be easily checked out that **x**_1_ + **x**_2_ − **x**_3_ = 0. It is seen that **x**_3_ = **x**_1_ + **x**_2_, and the set of vectors {**x**_1_, **x**_2_, **x**_3_} in ℝ^2^ is linearly dependent. As an immediate result, we can infer from this example that a linearly dependent set of vectors has a redundancy so that one of the vectors can be took away from the original vectors without affecting the span of the other vectors.

Considering what is described above, a significant advantage of removing linearly dependent features from the original set of features is that it helps to achieve one of the main goals of feature selection that is minimizing the redundancy. On the other hand, every basis for the feature space possesses two major characteristics. Firstly, it spans the feature space, and secondly, all the features included in a basis are linearly independent. Such a basis reduces the feature space so that the redundancy among the features decreases. It is because the reduced features are linearly independent as elements of the basis. based on this, the role that a basis for the feature space plays in learning tasks is investigated in more depth in this paper and this investigation involves the theoretical foundation as well as some practical aspects.

### 2.2 An Algorithm to Build a Basis for the Features Space

In this section, we aim to describe how to find a proper basis for the feature space. The technique of Variance Score (VS) is an effective method that determines a discriminative feature as the one whose VS is a large value. This technique is very simple to implement and is commonly used for feature selection problems in unsupervised mode. It is known that the variance of data represents the level of its spread over a given domain. The variance maximization for the data that are reduced and transformed into a lower dimension leads to far more information extraction. Considering the remarkable ability of this method, our proposed techniques apply it to determine which features are the most discriminative to be included in the basis we are going to form. Such a basis is constructed as follows. First, computing every feature’s VS, the features are reordered in terms of their VS so that the feature with the highest VS comes first, and the other features are ordered in a designing manner after it. Next, the first feature, the one whose VS is the highest value, is considered as the first feature of the basis and the set including this single feature is extended to a basis of features by applying the basis extension algorithm to the other features ordered after it. Algorithm 1 provides a detailed description of this process.

#### Algorithm 1 Building a basis for the feature space.

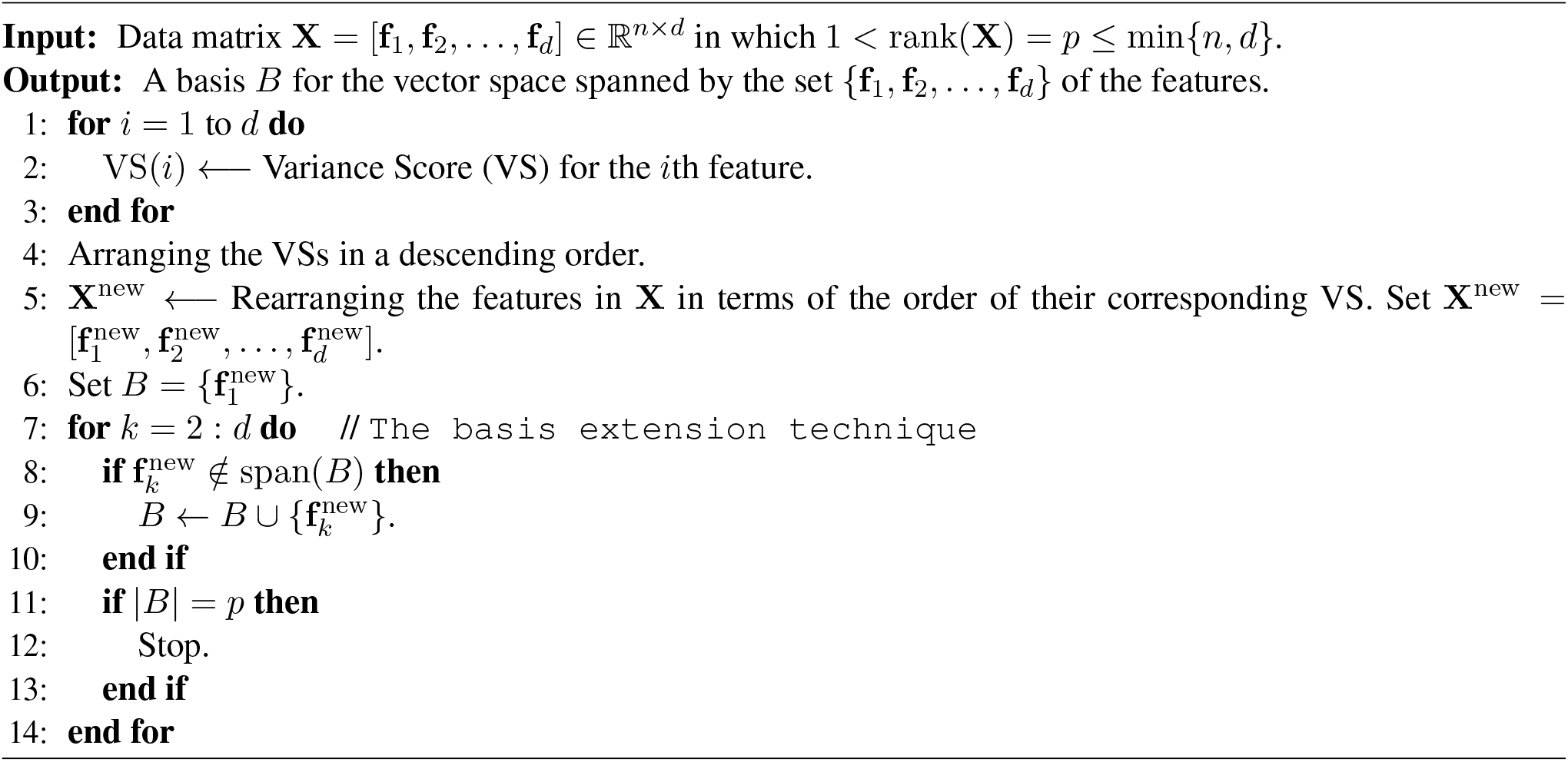

In Algorithm 1, the basis extension technique has been utilized to construct a basis for the feature space. In linear algebra, this technique is a standard commonly used way to produce a basis of a given vector space when the dimension of the space is finite [44]. For an *n* × *d* matrix **V** = [**v**_1_, …, **v**_*d*_], where 1 < rank(**V**) = *p* ≤ min {*n, d*}, it is assumed that 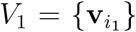 which 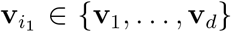 is a non-zero vector of size *n*. Therefore, the subset *V*_1_ is linearly independent. Now considering 1 < *p*, a vector 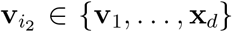 span(*V*_1_) can be found. It is evident that the extension of *V*_1_ defined as 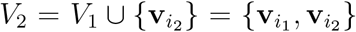 is a linearly independent subset. Now, continuing this process, a series *V*_3_, *V*_4_, …, *V*_*p*_ of subsets that are linearly independent extensions of *V*_1_ is obtained. Considering the fact that *V*_*p*_ is composed of *p* vectors, *V*_*p*_ is a basis of 𝒱 = span **v**_1_, …, **v**_*d*_. Algorithm 2 summarizes the procedure of the basis extension technique to construct a basis for span(**V**).

#### Algorithm 2 The basis extension technique [44].

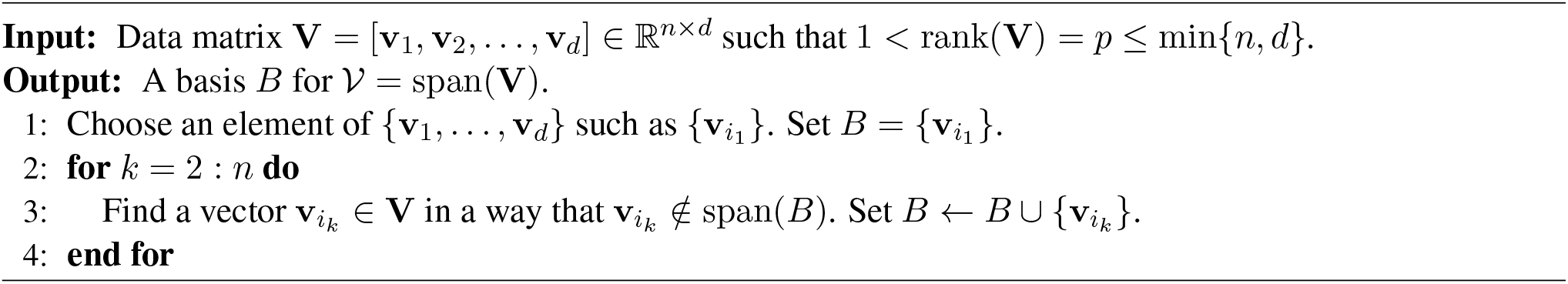

## 3 Proposed Frameworks

This section presents a set of novel and effective feature selection techniques rested on Problem (2) and the notion of a basis. In Section 3.1, an original unsupervised feature selection method is offered based on a basis of the feature space. In addition, Section 3.2 proposes a new unsupervised feature selection technique in which the redundancy among the features is minimized. Finally, Section 3.3 introduces another method for feature selection that preserves the intrinsic geometry of features.

### 3.1 The B-MFFS Method

A set of methods have been founded on the concept of matrix factorization, including Singular Value Decomposition (SVD) [45], Non-negative Matrix Factorization (NMF) [46] and Principal Components Analysis (PCA) [47]. Since such techniques were originally developed to handle feature extraction tasks, they are not always performed well enough when applied to feature selection problems. Especially, interpretation of the features determined by the mentioned methods in a subspace of lower dimension is not acceptable. One technique to deal with this problem was “Matrix Factorization criterion of Feature selection Subspace learning” (MFFS), proposed by Wang et al. [37]. In this approach, the matrix factorization was used for feature selection. This method was constructed using the problem presented in Eq. (6) and determines the optimal subset of features subject to an orthogonality condition imposed on its optimization problem.

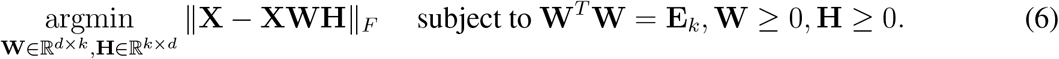

The current section aims to Considering the aforementioned reduced formspresent a novel method, B-MFFS, for feature selection problems that is rested on the mathematical notion of a basis. Particularly, the B-MFFS problem is established based on the idea of selecting a basis for the feature space as a representation of this space. This means that first, a basis is determined for the space of the original features. Then, to design an efficient and effective mechanism for feature selection, the B-MFFS algorithm represents the original feature space by the span of this basis and minimizes the distance between this span and the set of the selected features. In addition, making use of the *L*_2,1_–norm, the matrix of feature weights is regulated to guarantee the sparsity of feature selection [48].

Let *X* = {**f**_1_, **f**_2_, …, **f**_*d*_} be the set of the original features and 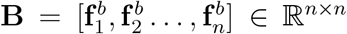 be a basis for the span of **X**. Since for a given submatrix **X**_*I*_ of **X** = [**f**_1_, **f**_2_, …, **f**_*d*_], there is a unique representation of **X**_*I*_ in the form of a linear combination of elements of **B**, a decomposition of **X**_*I*_ exists as follows: **X**_*I*_ = **BG**, where **G ∈** ℝ^*n*×*k*^. Considering the aforementioned reduced forms for **X** and **X**_*I*_, the feature selection problem for B-MFFS is formulated by Eq. (7).

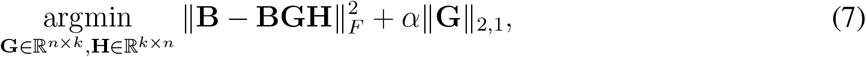

where *α* is a regularization parameter, and the *L*_2,1_–norm is imposed on the weight feature matrix **G** to guarantee that this matrix is sparse. Moreover, the term ∥**G**∥_2,1_ can be rewritten as:

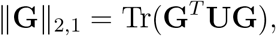

where **U** ∈ ℝ^*n*×*n*^ is a diagonal matrix with the diagonal elements

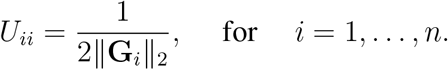

In order to find **G** and **H**, we make use of an alternating least squares iterative method in such a way that we will solve (7) with respect to one variable while the other one is taken to be fixed. For this purpose, we first assume that **H** ∈ ℝ^*k*×*n*^ is fixed. In this respect, it is supposed that

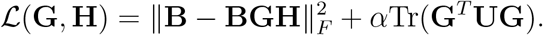

Now, by taking the partial derivative of ℒ with respect to **G**, we have

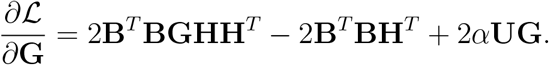

Set *∂*ℒ*/∂***G** = 0. It follows that the matrix **G** satisfies Eq. (8).

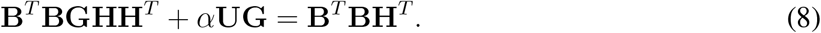

In the same manner, taking the partial derivative of ℒ with respect to **H** when **G ∈** ℝ^*n*×*k*^ is assumed to be fixed, we get

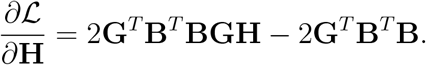

Again, putting *∂*ℒ*/∂***H** = 0, it is obtained that **H** satisfies Eq. (9).

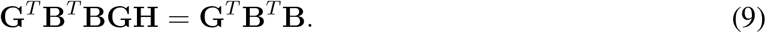

Algorithm 3 summarizes what has been discussed in this section.

#### Algorithm 3 The B-MFFS method.

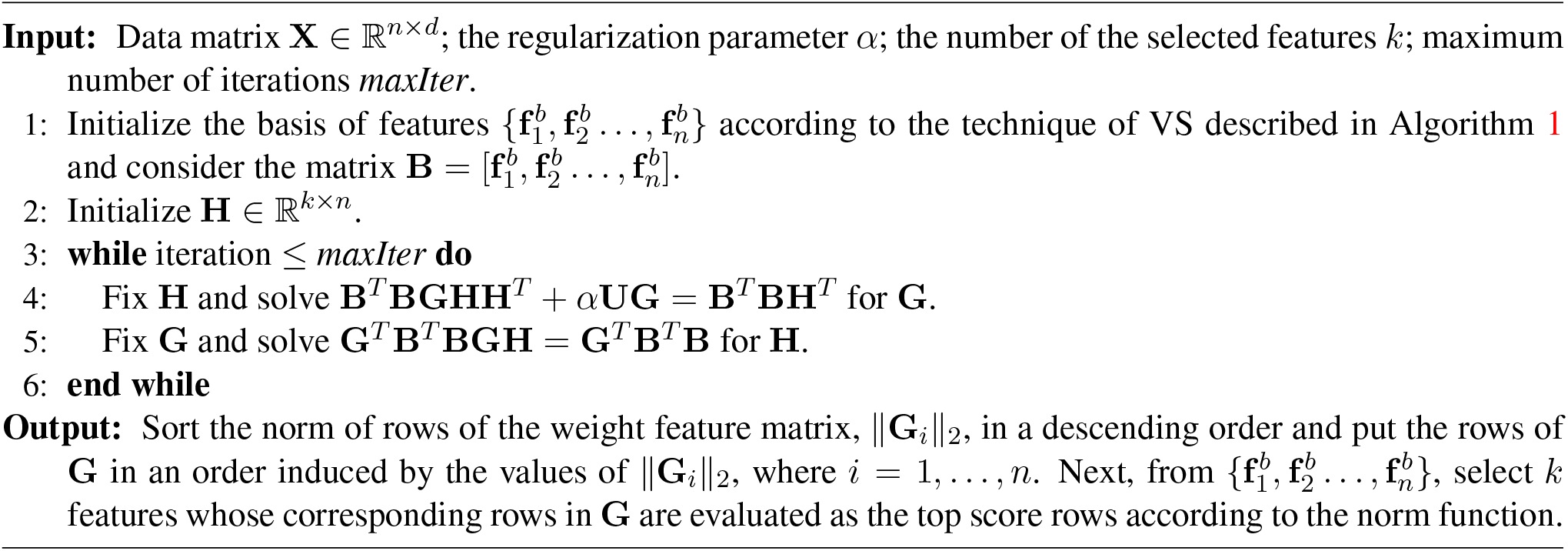

### 3.2 The B-MPMR Method

There are two major drawbacks when data of high-dimension are applied to machine learning methods. These include the significant increase in the storage costs and level of redundancy which negatively affect the learning process performance. To address these problems, a set of feature selection techniques have been developed. An example of such methods, that was successful in representing the original features effectively and reducing the features redundancy, is “feature selection based on Maximum Projection and Minimum Redundancy” (MPMR) [38], that can be formed as:

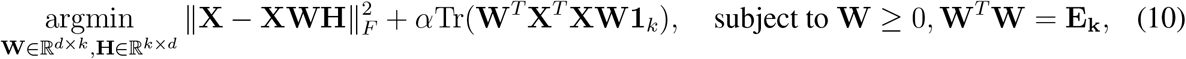

in which *α* is considered to achieve a proper trade-off between the redundancy rate and the approximation degree.

The current section introduces B-MPMR as an improvement of MPMR. The key idea in B-MPMR is to investigate that to what extent the features in a subset of the basis are independent of each other. Suppose that the basis 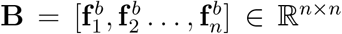 is as the one mentioned in Section 3.1. The B-MPMR technique addresses the redundancy problem by computing the rate of redundancy for each feature in the given feature subset and using the results to guide the process of selecting the features.

Given a submatrix of features, **X**_*I*_, there is a representation **X**_*I*_ = **BG** similar to the one described in Section 3.1. For **X**_*I*_, the rate of redundancy, red(*I*), is calculated by Eq. (11).

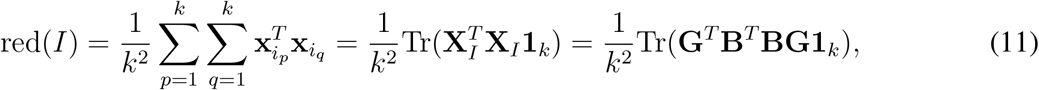

in which the matrix **1**_*k*_ ∈ ℝ^*k*×*k*^ is defined so that all its entries are one. In this regard, we can see that

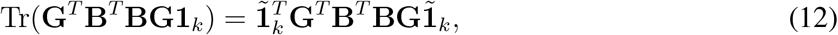

in which the vector 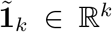 whose entries are all one. Considering the fact that **G**^*T*^ **B**^*T*^ **BG** is a symmetric positive semi-definite the matrix, it is deduced from Eq. (12) that Tr(**G**^*T*^ **B**^*T*^ **BG1**_*k*_) ≥ 0.

Regarding the redundancy problem discussed above, the feature selection problem of the B-MPMR method is formulated as follows:

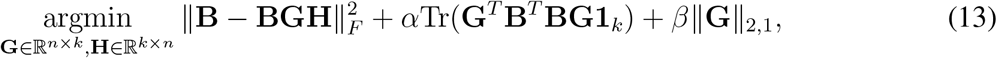

where *α* and *β* are two regularization parameters, and the *L*_2,1_–norm is imposed on the weight feature matrix **G** to assure the sparsity of this matrix.

To find **G** and **H**, the alternating least squares iterative method is employed again to solve (13) with respect to one variable while the other one is fixed. First, assume that **H ∈** ℝ^*k*×*n*^ is fixed. We define

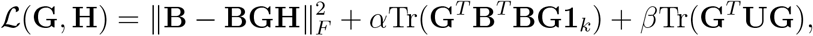

where **U ∈** ℝ^*n*×*n*^ is a diagonal matrix with the diagonal elements 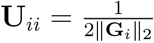 for *i* = 1, …, *n*. Now, by taking the partial derivative of ℒ with respect to **G**, we have

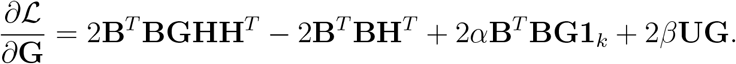

Next, assuming that *∂*ℒ*/∂***G** = 0, we get

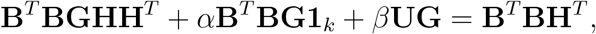

which can be expressed as

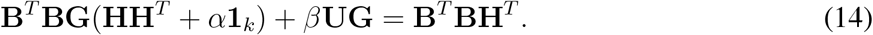

In the same manner, assuming that **G ∈** ℝ^*n*×*k*^ is fixed and taking the partial derivative of ℒ with respect to **H**, we have

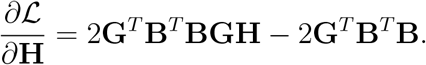

Setting *∂*ℒ/*∂***H** = 0, it follows that the matrix **H** satisfies Eq. (15).

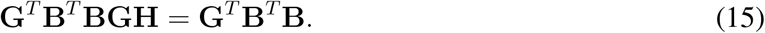

Algorithm 4 describes the steps of B-MRPR in brief.

### 3.3 The B-SGFS Method

The majority of former approaches consider the original space of features as the domain for the feature selection process. They also assume that there is a smooth manifold on which the original data are distributed [49, 50, 51]. For instance, the Subspace learning-based Graph regularized Feature Selection (SGFS) method [39] has been recently developed by including the notion of graph regularization in the objective function of a former feature selection method. This technique is formulated as:

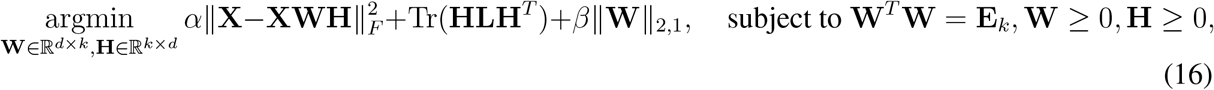

#### Algorithm 4 The B-MPMR method.

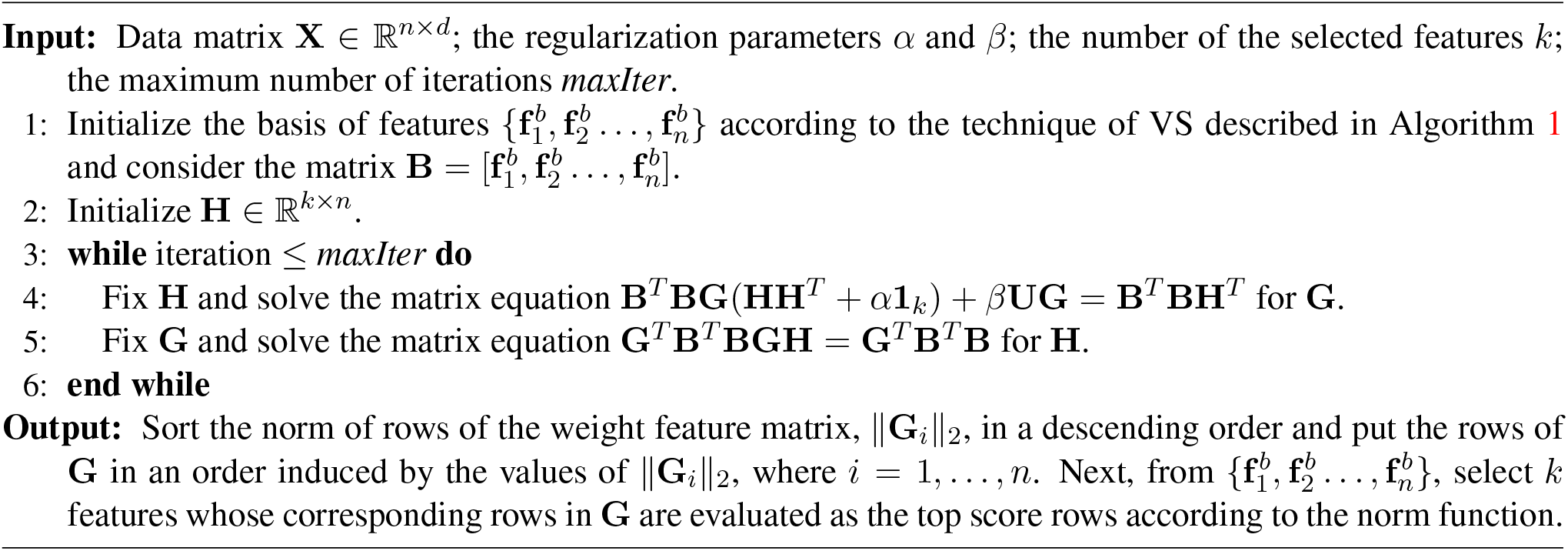

in which *α* and *β* are nonnegative balance parameters, and **L ∈** ℝ^*d*×*d*^ is the feature Laplacian matrix. In Eq. (16), the term Tr(**HLH**^*T*^) handles the geometric structure preservation task for the feature manifold. Moreover, to guarantee that the matrix of feature selection is sparse, this matrix is constrained by the *L*_2,1_–norm in the objective function of SGFS.

In this section, the B-SGFS method is presented as a technique to select the most discriminative features from a basis of the original feature space. This methodology also preserve the geometric structure of features associated with the basis manifold. This method is different from the SGFS method in terms of two properties including geometric structure preservation and feature manifold reduction. The main idea of our proposed B-SGFS method is to reduce the original feature graph through introducing a basis for the original feature space within which the selected feature subset is chosen. In other words, the reduced graph includes only the elements of the basis determined by the suggested method as its vertices. Instead of the original feature graph, the reduced graph can be applied as a model of the feature manifold. From a geometric point of view, this issue can be interpreted as follows. The manifold of features is represented by one of its sub-manifolds. This sub-manifold is determined by the basis that the proposed method introduces and it is composed of all the features included in that basis. Considering only the sub-manifold associated with a basis of features allows us to access the information about the whole feature manifold as well.

Let 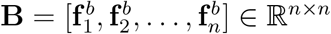 be a basis of features. Assuming that the basis of features exists on a sub-manifold of the original manifold of features, the similarity of the *i*th and the *j*th basis features for *i, j* ∈ {1, 2, …, *n*}, is denoted by 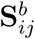 and is defined as:

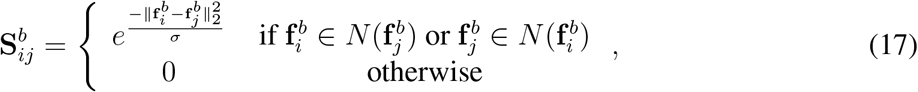

where the set 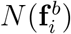 is the *k*-nearest neighbor corresponding to 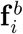, and *s* denotes the Gaussian parameter. In addition, the feature Laplacian matrix **L**^*b*^ corresponding to the basis of features is defined as **L**^*b*^ = **D**^*b*^ − **S**^*b*^ in which **D**^*b*^ is an *n* × *n* diagonal matrix with the diagonal entries 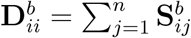. Suppose that the coefficient matrix is **H** = [**h**_1_, **h**_2_, …, **h**_*n*_]. The problem regarding the preservation of the geometric structure of features associated with the basis manifold is presented as:

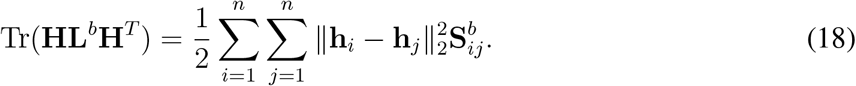

Considering Eq. (18), in case the level of the similarity between the *i*th and the *j*th features in the basis is sufficiently high, the similarity between the vectors **h**_*i*_ and **h**_*j*_ are also considerably strong. It means that the similarity of the feature vectors in the original manifold implies the similarity of the corresponding vectors in the sub-manifold determined by the basis. This representation leads to a substantial reduction of the feature manifold so that the local geometry of features is preserved while the size of the graph is enormously reduced. This issue enhances the performance of our method and reduces the computational costs significantly.

Considering the ultimate goal of an unsupervised feature selection to search a low-dimensional feature subset that best characterizes the original feature space, we define the optimization framework of the proposed method B-SGFS as follows:

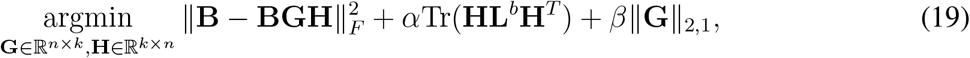

in which *α* and *β* are two regularization parameters, and the following principle fact has been utilized in (19): every feature subset of *X* can be expressed uniquely as a linear combination of the features associated with the basis *B*, and as an immediate consequence, it is deduced that there is a matrix **G ∈** ℝ^*n*×*k*^ such that **X**_*I*_ = **BG**.

In order to find **G** and **H**, we make use of an alternating least squares iterative method in such a way that we will solve (19) with respect to one variable while the other variable is assumed to be fixed. For this purpose, first, let **H** ∈ ℝ^*k*×*n*^ is taken to be fixed and suppose that

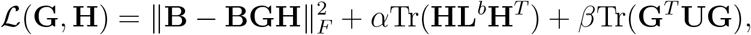

where **U** ∈ ℝ^*n*×*n*^ is a diagonal matrix with the diagonal elements 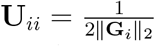 for *i* = 1, …, *n*. Now, by taking the partial derivative of ℒ with respect to **G**, we get

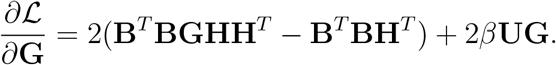

Next, we assume that *∂*ℒ*/∂***G** = 0. It follows that the matrix **G** satisfies Eq. (20).

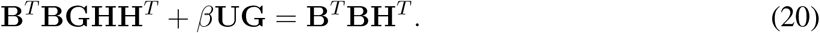

In the same manner, we assume that **G** ∈ ℝ^*n*×*k*^ is fixed and taking the partial derivative of ℒ with respect to **H**, we have

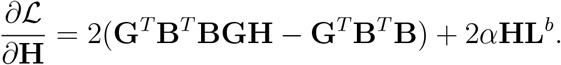

Now, suppose that *∂*ℒ*/∂***H** = 0. It results that the matrix **H** satisfies Eq. (21).

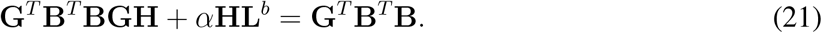

Algorithm 5 presents a summary of what discussed in this section.

#### Algorithm 5 The B-SGFS method.

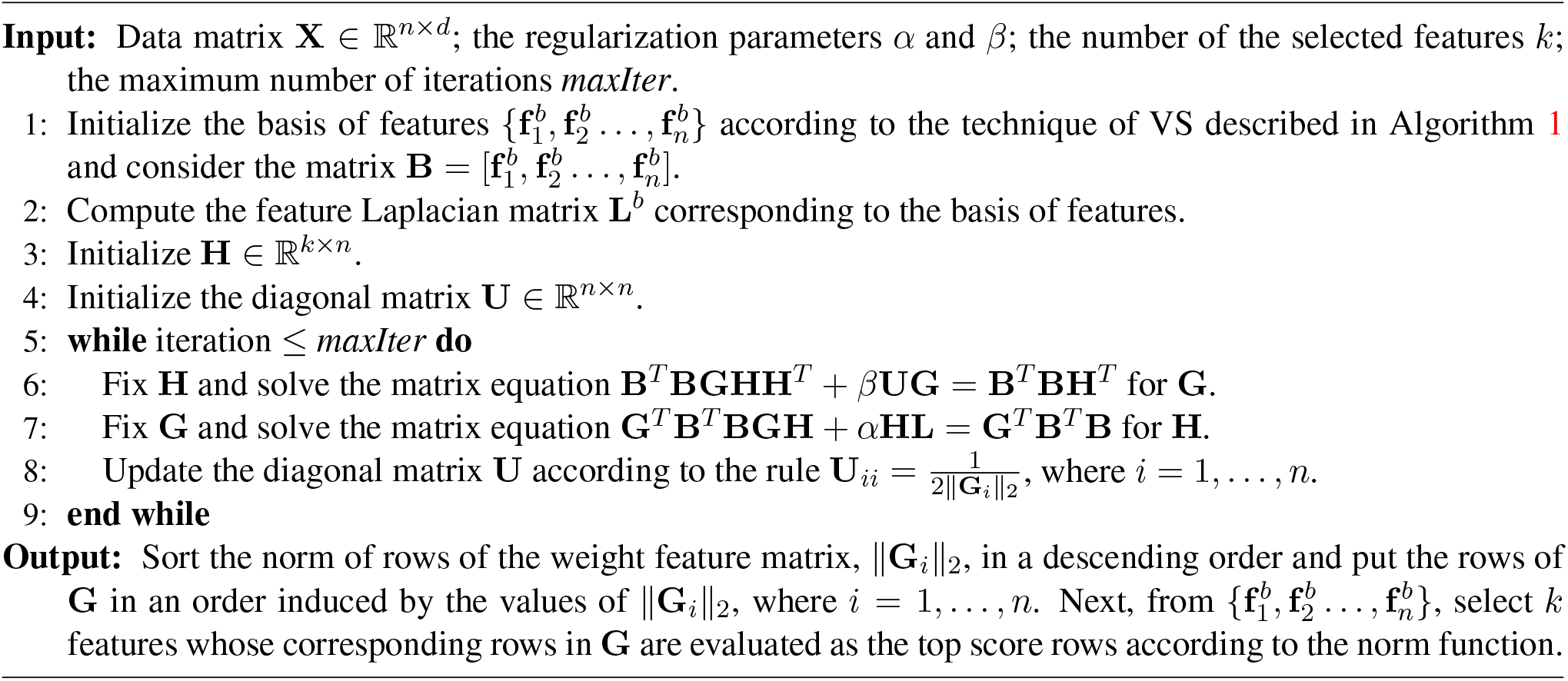

### 3.4 Solving the Matrix Equations in B-MFFS, B-MPMR and B-SGFS

As it was discussed in the previous sections, the following matrix equations have been derived for the three proposed methods B-MFFS, B-MPMR and B-SGFS.

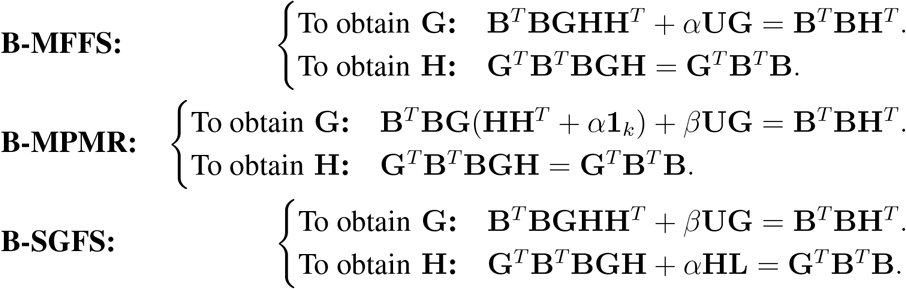

Since the coefficient matrices described in the above matrix equations are symmetric positive semidefinite, according to the results mentioned in [52], we can employ some reliable techniques such as the global conjugate gradient method to solve such matrix equations. In Appendix A, the details of the global conjugate gradient method will be explained. For more information about this method as well as the global version of this method, see [52, 53, 54] and references therein.

### 3.5 Comparison Between Our Proposed Methods and Corresponding Conventional Methods

One of the main advantages of using the basis of features in the frameworks presented in this study is that it significantly reduces the volume of calculations, and consequently, the time required in the process of reducing the high-dimensional space. Basically, the original formula used in the feature selection problem, presented in some state-of-the-art methods MFFS, MPMR and SGFS, is defined as:

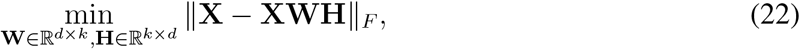

where **X** = [**f**_1_, **f**_2_, …, **f**_*d*_] ∈ ℝ^*n*×*d*^ is the data matrix, in which *n* and *d* indicates the number of samples and that of features, respectively. It should be here noted that in the high-dimensional datasets such as those examined in our experiments, the value of *d* can be very high. Furthermore, as it is discussed in the literature of the methods MFFS, MPMR and SGFS, the computational cost of these methods is strongly dependent on the number of features (i.e., *d*), and the greater the number of features, the longer the execution time of these methods [37, 38, 39].

On the other hand, the feature selection problem used in the frameworks presented in this study, i.e., B-MFFS, B-MPMR and B-SGFS, is defined as:

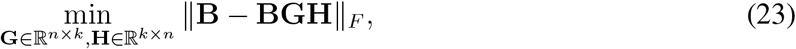

where 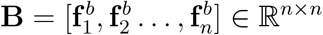 is a basis for the feature space generated by {**f**_1_, **f**_2_, …, **f**_*d*_}. Contrary to what is seen in Problem (22), the computational cost of our methods depends on the value of *n*, which is much smaller than *d*. In addition to this, a comparison between the size of the matrices **X** and **B** shows that the size of **B** can be much smaller than that of **X**. For this reason, it would be perfectly reasonable to expect that the time required for our proposed methods B-MFFS, B-MPMR and B-SGFS in the process of reducing the high-dimensional space to be much lower than the time required by the original methods MFFS, MPMR and SGFS.

In order to verify the theoretical results given here, the running time of all the proposed methods and their corresponding conventional versions are illustrated in Section 4.10. The numerical results indicate that the running time corresponding to all the proposed methods B-MFFS, B-MPMR and B-SGFS is much shorter compared to all the conventional methods MFFS, MPMR and SGFS. This means that applying the notion of basis to modify MFFS, MPMR and SGFS succeeds to enhance these methods in terms of their running time. As discussed above, the reason for such an improvement comes from the fact that representing the original data by a basis of the feature space reduces the consequent level of computations. Thererfor, he proposed methods present modifications of the corresponding conventional methods in terms of the amount of calculations and running time.

## 4 Computational Experimental Results

To validate the three proposed methods, B-MFFS, B-MPMR and B-SGFS, in terms of their performance, some experiments are conducted in this section on a number of well-known biological datasets. To analyze the results, two essential metrics are used. First, these metrics are described in brief. Following this, the results of the experiments regarding the efficiency and efficacy of the proposed techniques are discussed.

### 4.1 Datasets

n this paper, to evaluate the proposed methods, eight cancer gene expression datasets are used. A description of these datasets is presented in Table 1. Embryonal Tumors of Central Nervous System (CNS) dataset comprises tumor samples gene expression data from patients with different types of brain tumors. This dataset provides the gene expression profiles and clinical data on the progression of the disease and therapeutic responses. This dataset was used to predict the disease outcomes based on molecular signatures [55]. The Colon cancer dataset that we have used has gene expression profiles of the tumor and normal colon samples (40 tumor tissue samples and 22 normal colon tissue). This dataset has been used to detect gene expression signatures and patterns differentiating colon tissue types in normal conditions and neoplasia[56]. The dataset of Diffuse Large B-cell Lymphomas (DL-BCL) has the gene expression profiles of tumor samples from patients with DLBCL, which is the most common form of non-Hodgkin’s lymphoma [57]. Glioblastoma Multiforme (GBM) dataset includes gene expression profiles of samples from GBM tumors with the clinical characteristics of the patients in terms of outcome [58]. TOX-171 dataset is a toxicology-based gene expression set with integrated biological phenotypes and characteristics [28]. The Lung Cancer and Prostate Cancer datasets are integrated gene expression profiles from lung and prostate tumors [59]. The Cancer Cell Line Encyclopedia (CCLE) has the gene expression data of common cancer cell lines used in cancer research and contains metrics of the therapeutic responses and mutational properties of their genomes [60].

**Table 1:** The information of eight datasets utilized in the computational experiments.

| Dataset | Abbreviation | # Samples | # Features | # Classes | Reference |
| --- | --- | --- | --- | --- | --- |
| Embryonal Tumors of CNS | CNS | 60 | 7129 | 2 | [55] |
| Colon Cancer | Colon | 62 | 2000 | 2 | [56] |
| Diffuse Large B-cell Lymphomas | DLBCL | 47 | 4026 | 2 | [57] |
| Glioblastoma Multiforme | GBM | 85 | 22283 | 2 | [58] |
| TOX-171 | TOX-171 | 171 | 5748 | 4 | [28] |
| Lung Cancer | Lung | 73 | 325 | 7 | [59] |
| Prostate Cancer | Prostate | 102 | 5966 | 2 | [59] |
| Cancer Cell Line Encyclopedia | CCLE | 343 | 1900 | 2 | [60] |

### 4.2 Computational Experimental Setting

Before performing the experiments, the parameters included in the methods are set as follows. For all methods, the tuning of the regular parameters is carried out making use of the grid-search algorithm. This issue is a leading factor in drawing reasonable and fair comparisons. This way, we set *α* = 1 in MPMR and *ρ* = 10^8^ in MPMR and MFFS [37, 38]. In addition, the search domain is selected as {10^*i*^ | *i* = −8, −7, …, 7, 8} for the rest of the balancing parameters in all methods. Moreover, for all methods, the maximum number of iterations and the number of the selected features from each dataset are set to 30 and {10, 20, 30, …, 100}, respectively. For methods that preserve the geometric structure of features, {10^*i*^ |*i* = 0, 1, …, 5} is considered as the search domain for *σ*, and each neighbor size is selected to be 5 on each dataset. Employing the *k*-means technique, the performance of the process of clustering is examined subject to the selected features, where for a given dataset, the true number of the classes is denoted by *c*. When the feature selection task is performed and is finished, the selected features are used to guide the *k*-means algorithm to carry out the samples clustering task. Since the clustering performance of the *k*-means algorithm relies upon initialization, in the experiments, the clustering process is repeated 20 times where diverse initializations take place at random. Then, the average clustering results are reported.

### 4.3 Evaluation Metrics for Comparison

In order to carry out proper evaluation of the proposed and benchmark algorithms, two commonly used performance evaluation metrics for unsupervised methods, including Clustering Accuracy (ACC) and Normalized Mutual Information (NMI), are employed. Notice that a greater value of ACC/NMI corresponding to a method indicates that the method under consideration achieves a better performance.

The metric ACC is formulated by Eq. (24).

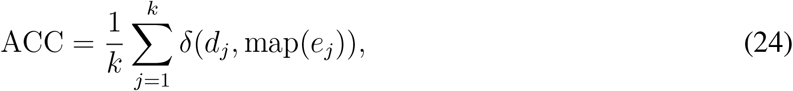

in which *k* denotes the total number of samples, *d*_*j*_ and *e*_*j*_ symbolize respectively the ground truth label for the sample *x*_*i*_ and the clustering label, map(.) represents an optimal mapping function that handles the task of matching the labels predicted by the given feature selection method to the real ones, and the indicator function *δ*(*a, b*) is defined by Eq. (25).

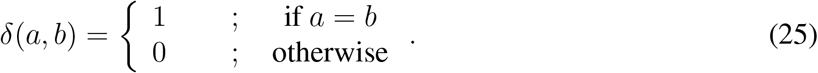

Let *W* and *Z* be random variables. The metric NMI is given by Eq. (26).

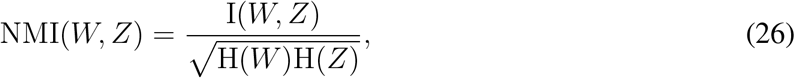

where I(*W, Z*) indicates the mutual information between *W* and *Z*, and H(.) is the entropy function.

### 4.4 Methods and Modalities

To conduct the experiments, we make comparisons between the proposed techniques and a number of benchmark methods including the Baseline, Variance Score (VS) [61], Laplacian Score (LS) [62], MFFS [37], MPMR [38] and SGFS [39]. In Baseline, all original features are used. Variance Score computes every feature’s variance, and determines the optimal features as the ones whose variance is the largest compared to the other features. In Laplacian Score, the intrinsic geometry of data is taken into consideration. The last three methods, MFFS, MPMR and SGFS, were also described in Section 3.

### 4.5 Results and Discussion

For comparisons of the proposed methods with other techniques, six various state-of-the-art methods were examined in terms of their performance on seven benchmark datasets. To this aim, ACC and NMI were computed for the selected methods. The results of these experiments are presented in Tables 2 and 3 in which the first and the second best results are marked in boldface and are underlined, respectively.

**Table 2:** Clustering accuracy (ACC±STD%) results for different feature selection methods on seven datasets. The numbers in parentheses are the numbers of selected features corresponding to the best clustering results. The best results in each row are highlighted in bold and the second best results are underlined. Note that the higher ACC values are, the better the clustering performance is.

| Dataset | Baseline | VS | LS | MFFS | MPMR | SGFS | B-MFFS | B-MPMR | B-SGFS |
| --- | --- | --- | --- | --- | --- | --- | --- | --- | --- |
| CNS | 53.33 ± 1.45 | 56.66 ± 0.04<br>(70) | 58.66 ± 1.02<br>(80) | 61.75 ± 2.78<br>(50) | 61.66 ± 0.00<br>(70) | 63.12 ± 0.00<br>(60) | 62.33 ± 2.05<br>(10) | <u>63.25 ± 0.37</u><br>(10) | **64.25 ± 3.35**<br>(20) |
| Colon | 78.54 ± 6.95 | 58.06 ± 1.14<br>(20) | 60.00 ± 9.38<br>(90) | 74.19 ± 0.29<br>(10) | 74.21 ± 2.91<br>(90) | 77.41 ± 1.45<br>(40) | 80.64 ± 0.03<br>(10) | <u>84.91 ± 2.52</u><br>(10) | **85.48 ± 0.00**<br>(10) |
| DLBCL | 59.57 ± 1.23 | 65.95 ± 0.01<br>(10) | 57.44 ± 0.72<br>(40) | 85.10 ± 0.14<br>(40) | 87.23 ± 0.14<br>(70) | 80.85 ± 0.43<br>(40) | **95.74 ± 0.29**<br>(10) | 87.23 ± 0.29<br>(10) | <u>90.85 ± 2.85</u><br>(10) |
| GBM | 67.05 ± 0.01 | 68.23 ± 0.00<br>(40) | 69.41 ± 0.00<br>(10) | 70.58 ± 0.00<br>(20) | 70.58 ± 0.00<br>(10) | 72.23 ± 0.00<br>(50) | <u>75.29 ± 0.00</u><br>(10) | <u>75.29 ± 1.45</u><br>(10) | **76.47 ± 0.26**<br>(10) |
| TOX-171 | 41.25 ± 0.72 | 42.13 ± 0.99<br>(20) | 40.35 ± 0.72<br>(10) | 42.13 ± 0.13<br>(90) | 41.49 ± 1.65<br>(60) | 38.26 ± 0.87<br>(100) | **45.08 ± 0.54**<br>(10) | <u>44.82 ± 0.91</u><br>(40) | 44.44 ± 0.72<br>(20) |
| Lung | 76.78 ± 5.03 | 59.58 ± 5.68<br>(100) | 65.82 ± 3.72<br>(100) | 77.80 ± 4.6130<br>(30) | 79.58 ± 6.57<br>(100) | 80.75 ± 5.45<br>(90) | 81.16 ± 4.96<br>(50) | **82.46 ± 5.83**<br>(50) | <u>82.05 ± 6.91</u><br>(50) |
| Prostate | 57.84 ± 0.00 | 65.68 ± 0.06<br>(10) | 60.78 ± 0.01<br>(10) | 61.76 ± 0.00<br>(50) | 62.84 ± 1.38<br>(60) | 63.72 ± 1.45<br>(10) | <u>69.60 ± 0.02</u><br>(10) | **69.72 ± 0.01**<br>(10) | 67.64 ± 0.00<br>(10) |

**Table 3:** Normalized mutual information (NMI±STD%) results for different feature selection methods on seven datasets. The numbers in parentheses are the number of selected features corresponding to the best clustering results. The best results in each row are highlighted in bold and the second best results are underlined. Note that the higher NMI values are, the better the clustering performance is.

| Dataset | Baseline | VS | LS | MFFS | MPMR | SGFS | B-MFFS | B-MPMR | B-SGFS |
| --- | --- | --- | --- | --- | --- | --- | --- | --- | --- |
| CNS | 1.17 ± 0.00 | 2.25 ± 0.45<br>(70) | 2.44 ± 0.45<br>(90) | 2.93 ± 0.45<br>(100) | 1.60 ± 0.80<br>(30) | **8.79 ± 0.00**<br>(30) | 3.91 ± 0.59<br>(10) | 3.61 ± 0.31<br>(30) | <u>4.26 ± 0.31</u><br>(20) |
| Colon | 27.35 ± 9.96 | 2.96 ± 1.63<br>(10) | 7.27 ± 1.23<br>(100) | 24.30 ± 0.14<br>(100) | 21.08 ± 0.54<br>(40) | 30.03 ± 0.00<br>(10) | 33.03 ± 0.01<br>(10) | <u>36.67 ± 5.43</u><br>(10) | **37.89 ± 1.82**<br>(10) |
| DLBCL | 2.93 ± 1.09 | 7.46 ± 1.13<br>(10) | 2.04 ± 0.45<br>(40) | 39.45 ± 0.00<br>(40) | 48.46 ± 0.72<br>(70) | 30.54 ± 0.72<br>(40) | **78.80 ± 0.00**<br>(10) | 48.46 ± 0.07<br>(10) | <u>58.06 ± 6.04</u><br>(10) |
| GBM | 21.74 ± 0.03 | 6.61 ± 1.82<br>(40) | 5.52 ± 2.75<br>(60) | 7.10 ± 0.18<br>(30) | 7.10 ± 0.82<br>(10) | 8.75 ± 0.00<br>(60) | 23.91 ± 0.36<br>(10) | <u>25.28 ± 1.09</u><br>(10) | **31.61 ± 0.27**<br>(10) |
| TOX-171 | 13.54 ± 0.23 | 12.27 ± 0.65<br>(20) | 11.68 ± 1.97<br>(90) | 12.82 ± 1.52<br>(80) | 16.22 ± 1.89<br>(60) | 10.41 ± 1.02<br>(100) | <u>17.39 ± 0.89</u><br>(50) | 16.83 ± 1.03<br>(10) | **18.51 ± 0.01**<br>(20) |
| Lung | 73.13 ± 2.52 | 56.36 ± 3.32<br>(100) | 62.85 ± 2.16<br>(100) | 73.22 ± 4.43<br>(60) | 73.67 ± 4.49<br>(100) | 75.61 ± 4.38<br>(100) | 76.41 ± 3.61<br>(50) | <u>77.29 ± 4.50</u><br>(50) | **79.01 ± 5.33**<br>(50) |
| Prostate | 1.85 ± 0.03 | 7.25 ± 0.00<br>(10) | 5.28 ± 0.06<br>(10) | 5.29 ± 0.03<br>(50) | 7.74 ± 1.28<br>(60) | 7.45 ± 0.91<br>(10) | <u>13.43 ± 0.79</u><br>(10) | **16.03 ± 0.27**<br>(10) | 10.21 ± 0.22<br>(10) |

According to Table 2, on all the datasets, the first and the second best values of ACC were achieved by the three proposed methods, B-SGFS, B-MPMR and B-MFFS. In addition, B-MFFS outperformed all the conventional methods in terms of ACC. With the exception of the ACC value for MPMR which equals that value for B-MPMR on DLBCL, B-MPMR gave a better performance compared to the conventional methods. It is also clear that B-SGFS showed a better performance compared to the state-of-the-art methods except for SGFS on CNS. In this case the difference between the ACC value of the two methods is very small. A key aspect of the proposed methods is the lower number of the selected features compared to that number corresponding to the conventional methods. It is clear from Table 2 that this number, presented in parentheses, is far smaller for our three methods than that number for the other methods in most cases which is a big advantage. Figure 2 illustrates the bar chart of ACC values for all the methods indicated in Table 2 on the seven benchmark datasets presented in Table 1.

**Figure 2:**
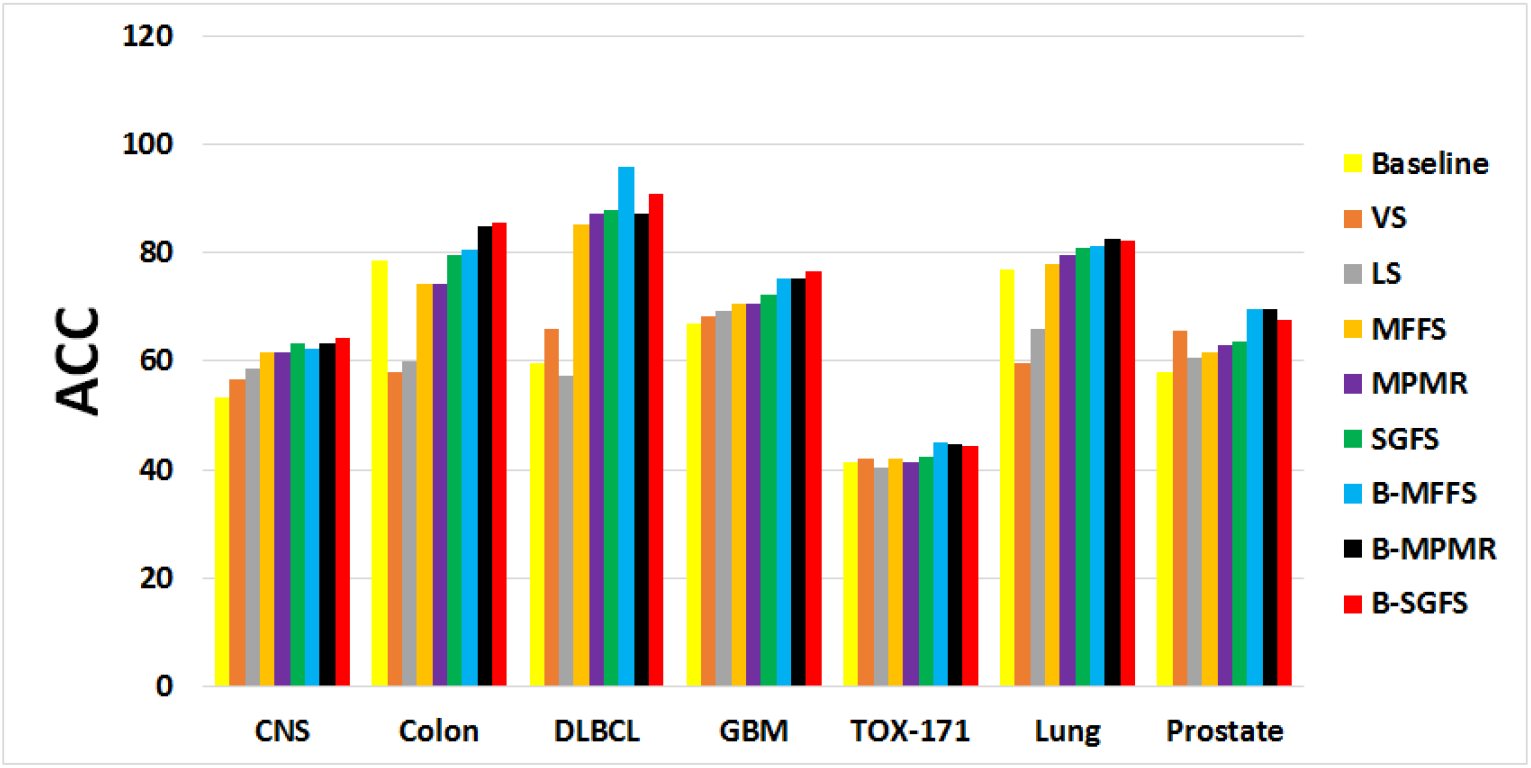
The average results of the ACC metric on seven datasets. The *x*-axis indicates the different datasets, and the *y*-axis represents the values of the ACC results obtained for each feature selection method. Note that the higher ACC values are, the better the clustering performance is.

The blue, black and red bars that correspond to B-SGFS, B-MPMR and B-MFFS, respectively, show the superiority of our proposed methods over all the other ones except for two mentioned cases.

Moving to Table 3, it is clear that B-MFFS, B-MPMR and B-SGFS are infinitely preferable in terms of NMI than the conventional methods on all the datasets used in this paper with the exception of two cases. The first exception occurred for SGFS on CNS, where this algorithm performed better compared with all our three methods, and the second one was raised on DLBCL in which the values of NMI for B-MPMR and MPMR were equal. In all the other cases, the first and second best results were obtained by the proposed methods. Again, the number of the selected features can be considered as an outstanding achievement of our methods since in almost all cases, these algorithms selected lower number of the features in compared to the conventional methods. In Figure 3, the bar chart of the values of NMI corresponding to the proposed and conventional methods on the seven benchmark datasets are shown. It can be seen from Figure 3 that B-MFFS, B-MPMR and B-SGFS, illustrated by blue, black and red bars, respectively, outperformed the conventional methods with two aforementioned exceptions.

**Figure 3:**
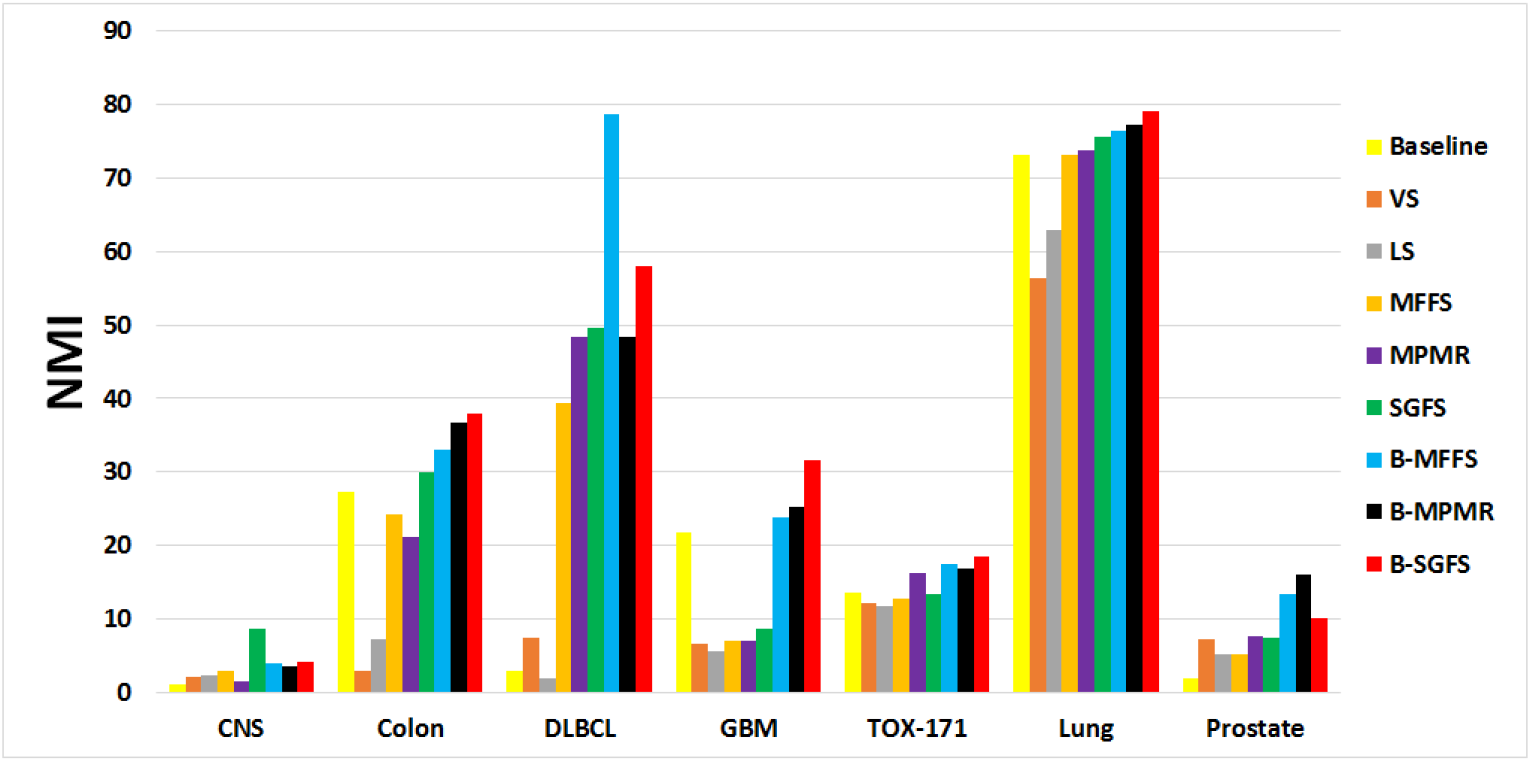
The average results of the NMI metric on seven datasets. The *x*-axis indicates the different datasets, and the *y*-axis represents the values of the NMI results obtained for each feature selection method. Note that the higher NMI values are, the better the clustering performance is.

Furthermore, Figure 4 presents the bar charts of both the ACC and NMI values of all the conventional and proposed algorithms averaged over all the benchmark datasets. This figure shows that for all the three proposed methods, the average values of ACC and NMI, illustrated by the gray and red bars, respectively, are higher than that values corresponding to the conventional methods. This means that overall, the B-MFFS, B-MPMR and B-SGFS algorithms show a stronger performance in terms of ACC and NMI compared to the benchmark methods.

**Figure 4:**
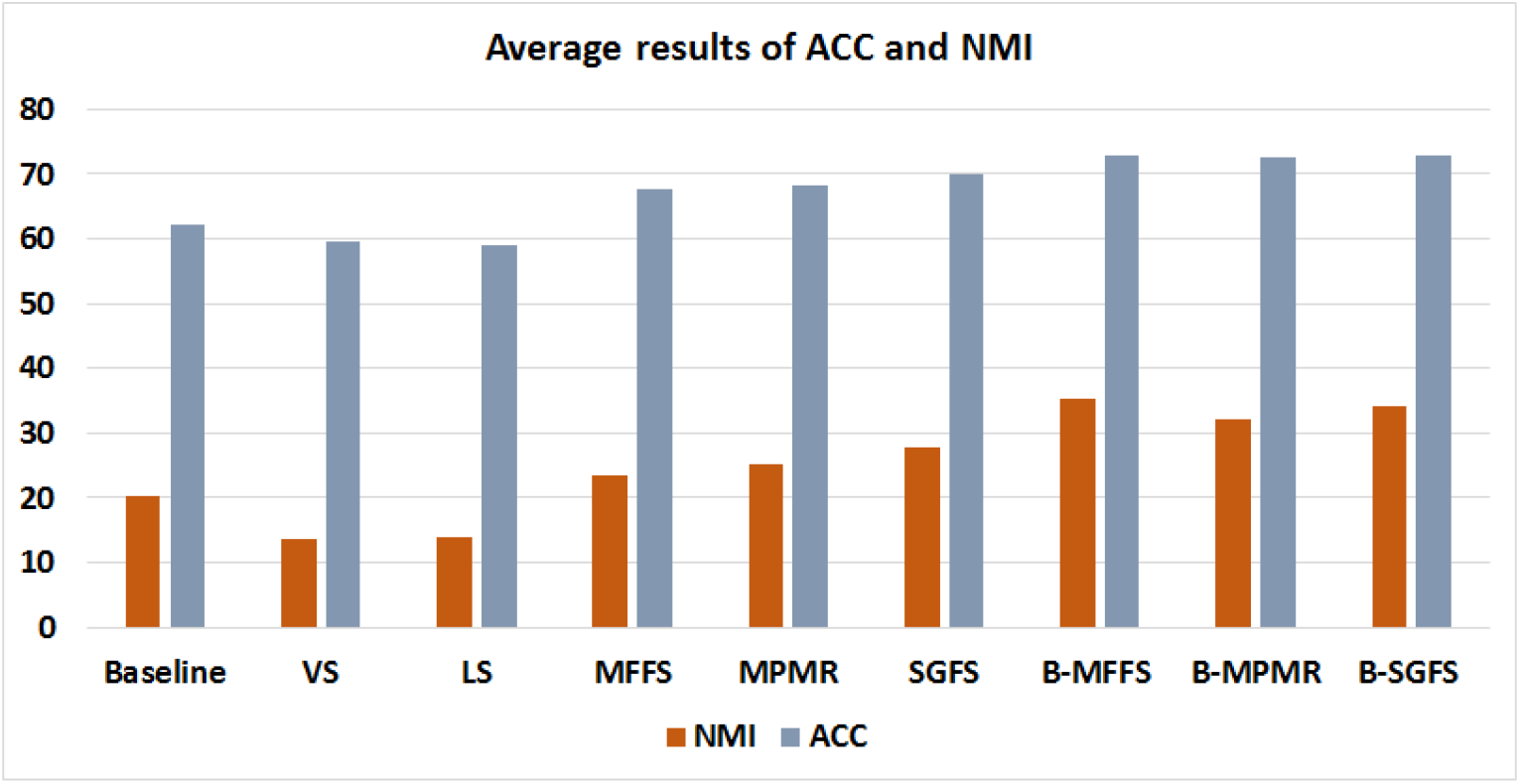
The average values of the metrics ACC and NMI. The *x*-axis indicates the different feature selection methods, and the *y*-axis represents the obtained values of the metrics. Note that the higher ACC and NMI values are, the better the clustering performance is.

### 4.6 Comparison Between MFFS and B-MFFS

The data in Table 2 show that in terms of ACC, B-MFFS outperforms the MFFS on all the datasets. Those data also indicate that B-MFFS stands at the first and second ranks on two datasets, DLBCL and TOX-171, and on the dataset GBM, respectively. In addition, it is apparent from Table 2 that compared to MFFS, B-MFFS selects a lower number of features on all datasets except for Lung, where both numbers are equal. Based on the data given in Table 3, it is clear that B-MFFS is superior in terms of NMI to MFFS on all the datasets and comparing all the considered methods, its results achieves the first and the second ranks respectively on the dataset DLBCL and on the datasets TOX-171 and Lung. Moreover, in terms of NMI, the number of the features selected by B-MFFS is lower than that number corresponding to MFFS. In Figure 5, the bar charts of the values of ACC and NMI, relevant to MFFS and B-MFFS and averaged over all the benchmark datasets, are presented. As Figure 5 shows, the average values of both indices for B-MFFS, represented by blue bars, are greater than that values for B-MFFS, represented by red bars. Thus, the average performance of B-MFFS is better than that of its corresponding conventional method.

**Figure 5:**
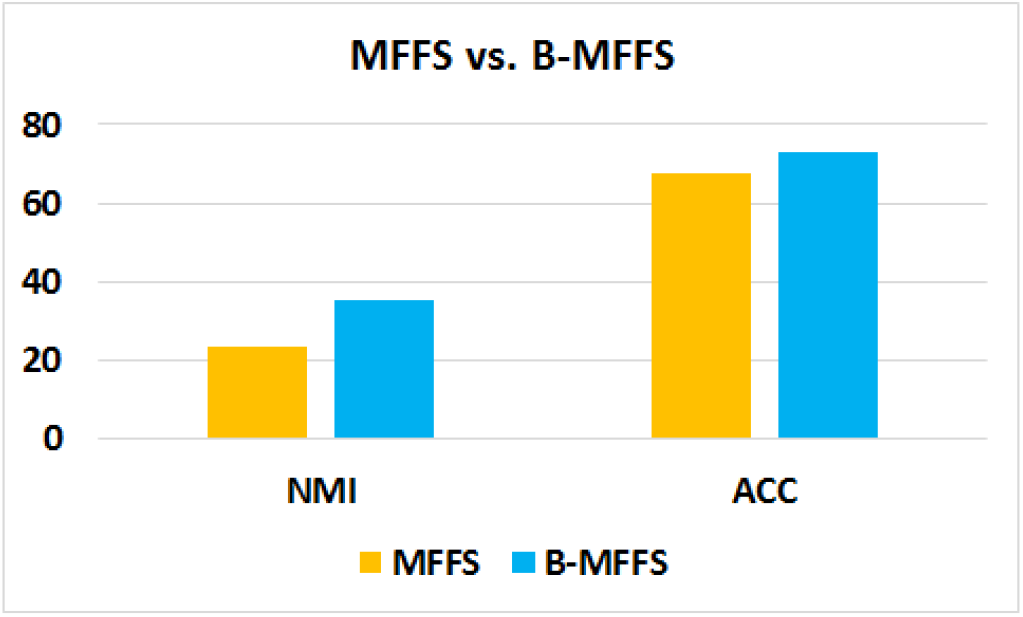
Comparison results of the MFFS and B-MFFS methods in respect of the average values of the metrics ACC and NMI.

### 4.7 Comparison Between MPMR and B-MPMR

To compare B-MPMR with its corresponding conventional method, the data in Tables 2 and 3 should be considered. It is obvious from these tables that in terms of both ACC and NMI, B-MPMR performs better than the MPMR over all the datasets except for DLBCL over which both the methods have the same performance. Furthermore, compared to all the methods considered in the experiments, B-MPMR stands at the first and the second ranks in terms of ACC respectively on two datasets Lung and Prostate and on the datasets CNS, Colon, GBM and TOX-171. In terms of NMI, B-MPMR also achieves the first and second best results on the dataset Prostate and on the datasets Colon, GBM and Lung, respectively. Moreover, it can be seen from Tables 2 and 3 that the number of the features selected by B-MPMR is lower compared to that number for MPMR over all the datasets with three exceptions including GBM in Table 2 and CNS and GBM in Table 3, where both methods selects the same number of features. The bar charts of the ACC and NMI values for MPMR and B-MPMR, averaged over all the benchmark datasets, are depicted in Figure 6. It is apparent from Figure 6 that the overall performance of B-MPMR is stronger in terms of ACC and NMI compared to that of MPMR since for B-MPMR, the ACC and NMI average values, that are illustrated by black bars, are greater compared with that values for MPMR, which are represented by purple bars.

**Figure 6:**
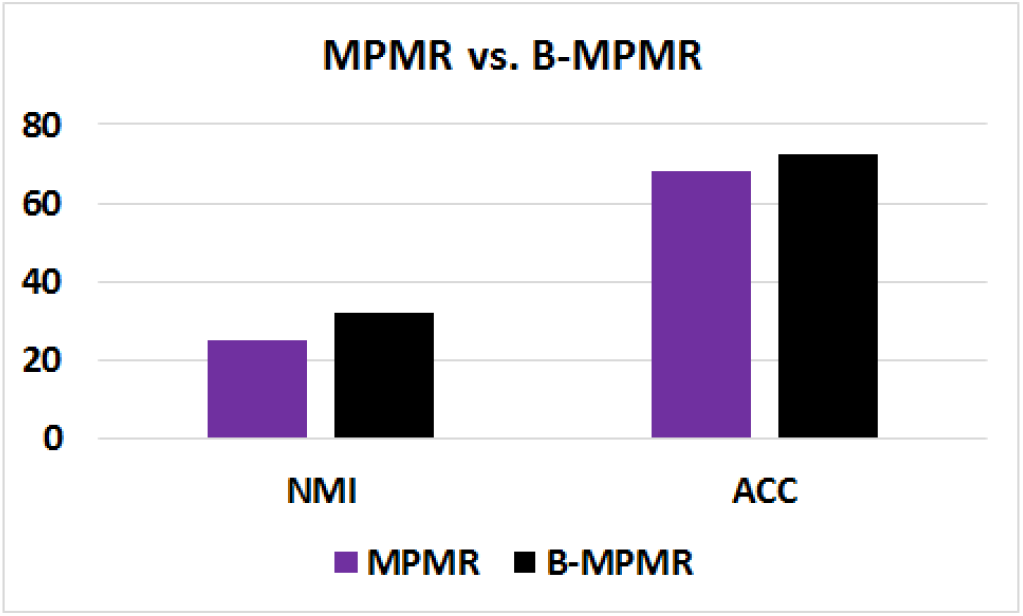
Comparison results of the MPMR and B-MPMR methods in respect of the average values of the metrics ACC and NMI.

### 4.8 Comparison Between SGFS and B-SGFS

As it can be seen from Tables 2 and 3, the values of ACC and NMI for B-SGFS is better than that of SGFS over all the datasets with the exception of CNS over which the NMI value for SGFS is greater than that of B-SGFS. In addition, compared to all the conventional methods and the two other proposed methods, B-SGFS holds the first and the second ranks in terms of ACC over the datasets CNS, Colon and GBM and over the datasets DLBCL and Lung, respectively. When considering NMI, the first and the second best results of B-SGFS is obtained over the datasets Colon, GBM, TOX-171 and Lung and on the datasets CNS and DLBCL, respectively. Furthermore, B-SGFS selects a smaller number of features in compared to SGFS over all the benchmark datasets except for Prostate in Table 2 and Colon and Prostate in Table 3, where the number of features selected by both methods are the same. Figure 7 illustrates the bar charts of the ACC and NMI values averaged over all the benchmark datasets corresponding to SGFS and B-SGFS. In this figure, the green bars represent the average value of SGFS and the red bars illustrate that of B-SGFS. From Figure 7, it is clear that B-SGFS achieves a better average performance in terms of ACC and NMI than does SGFS.

**Figure 7:**
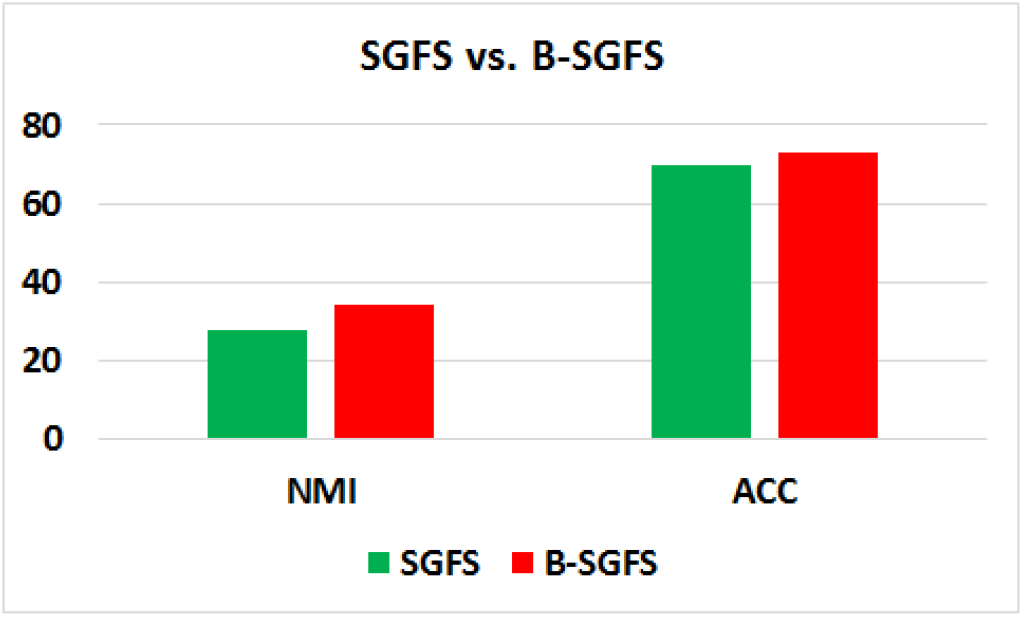
Comparison results of the SGFS and B-SGFS methods in respect of the average values of the metrics ACC and NMI.

### 4.9 Comparison Among the Proposed Methods

To compare the proposed methods with each other, first, we consider their values of ACC in Table 2. Based on these data, B-SGFS holds the first rank on datasets CNS, Colon and GBM. Over CNS and Colon, B-MPMR stands at the second rank and on GBM, both B-MFFS and B-MPMR achieves the second rank simultaneously. The B-MFFS algorithm obtains the best results on the datasets DL-BCL and TOX-171. The B-SGFS algorithm stands after it on DLBCL, while the B-MPMR algorithm produces the second best results on TOX-171. Finally, on the datasets Lung and Prostate, B-MPMR stands at the first rank. On Lung, B-SGFS holds the second rand while over Prostate, B-MFFS achieves the second best results. In terms of the number of the selected features, B-MFFS obtains the best results since it selects 10 features over all the benchmark datasets except for Lung over which it selects 50 features that is the same as the number of features the two other proposed methods select on this dataset. The B-MPMR algorithm selects 10 features over the rest of the datasets with the exception of TOX-171 over which it selects 40 features, while B-SGFS selects 20 features on CNS and TOX-171 and 10 feature on the rest of the datasets except for Lung.

Moving to Table 3, the NMI values are considered to make comparisons of our proposed methods. It is clear form this Table that with the exception of CNS on which SGFS produces the best results, for rest of the datasets, our methods obtains the best results. Even on CNS, B-SGFS holds the second best rank. This method also stands at the first rank over four datasets including Colon, GBM, TOX-171 and Lung. The B-MPMR method stands after it over Colon, GBM and Lung, while B-MFFS holds the second rank on TOX-171. The data in Table 3 also indicates that on the dataset Prostate, B-MPMR and B-MFFS achieve the first and the second best results, respectively, while over the dataset DLBCL, B-MFFS and B-SGFS produce respectively the first and the second best outcomes. Moreover, B-MPMR selects 30 and 50 features on CNS and Lung, respectively, and it selects 10 features over the rest of the datasets. In terms of the number of the selected features, B-MPMR perform better than the two other proposed methods. The B-MFFS algorithm selects 50 features on each of TOX-171 and Lung, while it selects 10 features over the rest of the datasets. The number of the features selected by B-SGFS is 20 on each of CNS and TOX-171, 50 on Lung and 10 over the rest of benchmark datasets.

### 4.10 Methods and Their Running Times

In Table 4, the running time of all the proposed and the conventional methods are illustrated on all the benchmark datasets and for the case of *k* = 40.

**Table 4:** The CPU time in seconds for B-MFFS, B-MPMR, and B-SGFS as well as for their corresponding conventional methods. Here, the number of the selected features is set to *k* = 40.

| Dataset | MFFS | MPMR | SGFS | B-MFFS | B-MPMR | B-SGFS |
| --- | --- | --- | --- | --- | --- | --- |
| **CNS** | 20.11 | 25.28 | 55.61 | 3.03 | 4.22 | 5.16 |
| **Colon** | 1.58 | 2.03 | 4.32 | 3.33 | 3.55 | 4.28 |
| **DLBCL** | 6.48 | 8.13 | 17.52 | 2.01 | 2.15 | 2.46 |
| **GBM** | 190.15 | 239.18 | > 1000 | 2.05 | 2.34 | 3.18 |
| **TOX-171** | 12.82 | 16.52 | 35.82 | 2.75 | 2.80 | 3.43 |
| **Lung** | 1.06 | 1.07 | 1.1 | 1.46 | 1.43 | 1.78 |
| **Prostate** | 14.07 | 18.34 | 39.68 | 2.35 | 2.41 | 3.32 |

Table 4 shows that the running time corresponding to all the proposed algorithms is much shorter compared to all the conventional methods with one exception including MFFS, MPMR, and SGFS over the Lung dataset. In specific, it is clear that for the case of GBM, the running time of all the three proposed methods is far less than that of MFFS, MPMR, and SGFS.

In addition, the CPU time in seconds, required to build the basis matrix corresponding to each dataset, is reported in Table 5. It can be seen from this table that building the basis matrix needs a very short time to be completed. In point of fact, since the size of the basis matrix is directly related to its rank, the higher the rank of the dataset is, the longer the CPU time will be.

**Table 5:** The CPU time in seconds required for building the basis matrix corresponding to each dataset.

|  | CNS | Colon | DLBCL | GBM | TOX-171 | Lung | Prostate | CCLE |
| --- | --- | --- | --- | --- | --- | --- | --- | --- |
| **Rank of dataset** | 60 | 62 | 47 | 85 | 171 | 73 | 102 | 343 |
| **CPU time** | 0.0222 | 0.0261 | 0.0174 | 0.0478 | 0.2133 | 0.0413 | 0.0598 | 1.8154 |

The most significant finding that Tables 4 and 5 provide is that all the proposed methods are faster than their corresponding conventional methods, MFFS, MPMR and SGFS, over all datasets (Lung is excepted). This means that applying the notion of basis to modify MFFS, MPMR and SGFS succeeds to enhance these methods in terms of their running time. The reason for such an improvement lies in the fact that representing the original data by a basis of the feature space reduces the consequent level of computations. Hence, the proposed methods present modifications of the corresponding conventional methods in terms of the amount of calculations and running time as well as improving their performance as mentioned earlier.

## 5 Application to the Extraction of Gene Signatures for Drug Responses in Cancer

Over the last recent years, dimensionality reduction techniques and feature selection methods have been applied to a variety of real-world problems in different fields, particularly to large molecular and clinical data including gene expression datasets [63, 64]. For instance, to study complex diseases such as cancer from a molecular perspective, it is required to reduce the dimension of the molecular data and to deal with the noise and redundancy included in such types of data. In this regard, a combination of molecular signature approaches with feature selection and reduction techniques can substantially help to provide new solutions.

In this section, the newly developed methods, B-MFFS, B-MPMR and B-SGFS, are incorporated into gene signature extraction to develop an analytical framework for cancer cell lines’ responses to Nilotinib treatment. In addition, the efficiency and productivity of these methods are evaluated using the CCLE dataset [60]. Furthermore, the proposed methods are compared to MFFS, MPMR and SGFS in terms of their performance.

### 5.1 Computational Experimental Metrics

To assess the proposed methods in terms of their classification performance and to corroborate their efficiency and efficacy, four types of evaluation metric including the classification accuracy (ACC), the true positive rate (TPR), the true negative rate (TNR), and the F1-score are employed. For a given dataset, these metrics are defined as:

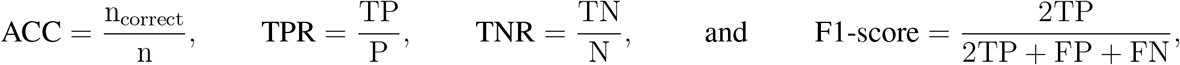

where *n* and *n*_correct_ indicate the number of the whole samples and the correct classified samples, respectively. Moreover, TP denotes the number of true positive samples, TN is the number of true negative samples, FP is the number of false positive samples, FN is the number of false negative samples, P indicates the number of positive samples and N is the number of negative samples.

The selected gene sets discovered by our proposed methods in each run have different sizes, specifically, the number of the selected genes is set to *k* = 2, 5, 10. It should be pointed out that when the feature selection process is completed, the methods used for feature selection are appraised in terms of their performance making use of the Support Vector Machine (SVM) classifier [65]. To draw a comparison between the proposed and the benchmark feature selection techniques, a 10-fold cross-validation (CV) process is applied such that the whole dataset is randomly splited into 10 groups in each CV process iteration. Then, nine of these groups are used to handle the classifier training phase and the remaining one is utilized to perform the testing phase. This process guarantees the generalization ability of the trained classifier to make accurate predictions of unseen samples. Moreover, the tuning process for setting the hyperparameters of the methods in the training phase is guided in each iteration by an internal five-fold CV grid search on the values presented in Subsection 4.2.

### 5.2 Results and Discussion

The experimental results regarding the performance of the proposed methods and its comparison with that of the benchmark methods are illustrated in Table 6 and Figure 8. It can be seen from Table 6 that the results of ACC, TNR and F1-score corresponding to the proposed methods B-MFFS, B-MPMR and B-SGFS show a better performance compared to that of the conventional methods MFFS, MPMR and SGFS. It is only in the case of TPR that the performance of the proposed methods is at a level the same as the conventional methods or is a bit lower than the performance of those techniques. Furthermore, in terms of ACC, TPR and TNR, the B-MFFS method outperforms B-SGFS and B-MPMR, while in terms of F1-score, B-SGFS performs better than the two other methods.

**Table 6:** Classification performance (ACC%, TPR%, TNR%, and F1-score%) results for different feature selection methods on the CCLE dataset. The number of the selected genes is set to *k* = 2, 5, 10, and the numbers in parentheses are the numbers of selected genes corresponding to the best classification results. The best results in each row are highlighted in bold and the second best results are underlined. Note that a higher ACC, TPR, TNR, and F-score value is indicative of a better classification performance.

| Dataset | MFFS | MPMR | SGFS | B-MFFS | B-MPMR | B-SGFS |
| --- | --- | --- | --- | --- | --- | --- |
| ACC | 72.68 (5) | 71.70 (2) | 72.59 (5) | **75.47 (2)** | 72.64 (2) | <u>80.19 (5)</u> |
| TPR | **98.00 (10)** | **98.00 (2)** | **98.00 (2)** | **98.00 (10)** | <u>97.35 (2)</u> | **98.00 (10)** |
| TNR | 20.13 (2) | 16.67 (5) | 12.32 (5) | **46.67 (2)** | 33.56 (2) | <u>43.33 (10)</u> |
| F1-score | 83.89 (5) | 83.47 (2) | 83.70 (5) | 89.37 (2) | <u>87.86 (2)</u> | **87.27 (5)** |

**Figure 8:**
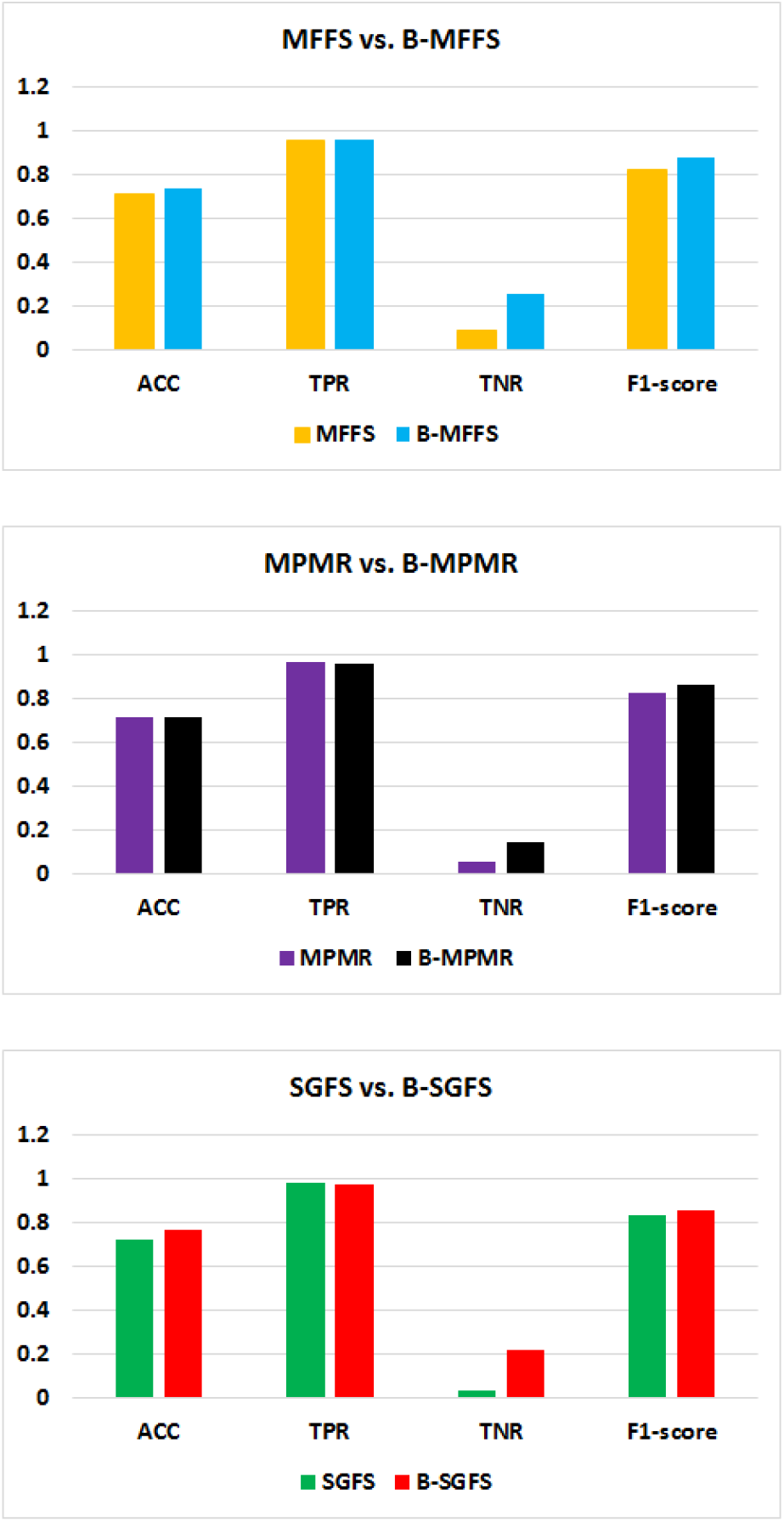
Results of comparison between “the MFFS and B-MFFS methods”, “the MPMR and B-MPMR methods”, and “the SGFS and B-SGFS methods” in respect of the average values of the metrics ACC, TPR, TNR, and F1-score.

Figure 8 presents the results of the comparisons between each of the proposed methods and its conventional technique in terms of the average values of ACC, TPR, TNR and F1-score. It is clear from this figure that in terms of ACC, MPMR and B-MPMR have the same performance while B-MFFS and B-SGFS perform better than their conventional methods. Considering TPR, the performance of B-MFFS, B-MPMR and B-SGFS is similar to that of MFFS, MPMR and SGFS, respectively. Finally, all the proposed methods outperform their corresponding conventional technique in terms of TNR and F1-score.

Overall, the outcomes of experiments demonstrate that the incorporation of the notion of a basis into MFFS, MPMR and SGFS is completely effective.

### 5.3 Verification of the Feature Selections in Therapeutic Responses to Nilotinib

Oxidative response is one of the pathways activated upon Tyrosine Kinase Inhibitors (TKI) treatment in cells [66, 67, 68]. We used the gene sets assigned to Oxidative Response in cells from Molecular Biology of The Cell (MBC) ontology [69]. To find a gene signature extracted from Oxidative Stress response pathway, we used the Cancer Cell Line Encyclopedia (CCLE) data on drug treatment [60]. The dataset we used contained IC_50_ for Nilotinib (a TKI inhibitor) treatment in 107 cancer cell lines IC_50_ ranges of 0.0029 to 7.98 *µM*) and their gene expression profiles. The aim was to find gene sets that can predict the cell lines’ responses to Nilotinib in terms of IC_50_ < 4*µM*. These gene sets were selected from oxidative stress pathway. The gene sets we discovered, had different sizes (gene numbers, *k* = 2, 5, 10). Our results demonstrate that applying B-MFFS and B-MPMR, we even can use a predictive 2-gene signatures. Table 7 shows which two genes are the most significant ones in each of these gene sets.

**Table 7:**
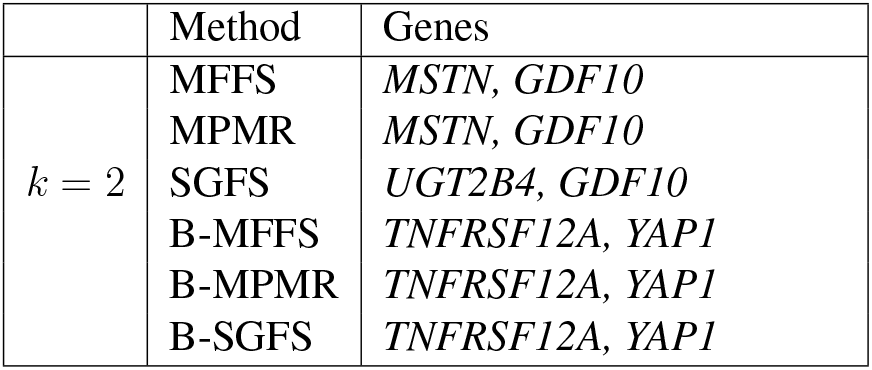
2-gene signatures selected by different feature selection methods.

Figure 9 illustrates how the proposed techniques, B-MFFS, B-MPMR and B-SGFS, and their corresponding conventional methods differentiate resistant cases from sensitive ones (using IC_50_ = 4*µM* as a threshold) with respect to two genes selected by each one of these methods as the selected features. It is apparent from Figure 9 that while MFFS, MPMR and SGFS are not able to clearly distinguish between the two categories of resistant and sensitive cases, the proposed methods carry out this task in a highly satisfying manner. All three B-MFFS, B-MPMR and B-SGFS techniques selected *TNFRSF12A* and *YAP1* while MFFS and MPMR selected *MSTN* and *GDF10* as the selected 2-gene signature. SGFS found *GDF10* and *UGT2B4* as selected features for a 2-gene signature. It has been shown that both *TNFRSF12A* and *YAP1* are involved in TKI treatment responsiveness in cancer [70, 71, 72, 73].

**Figure 9:**
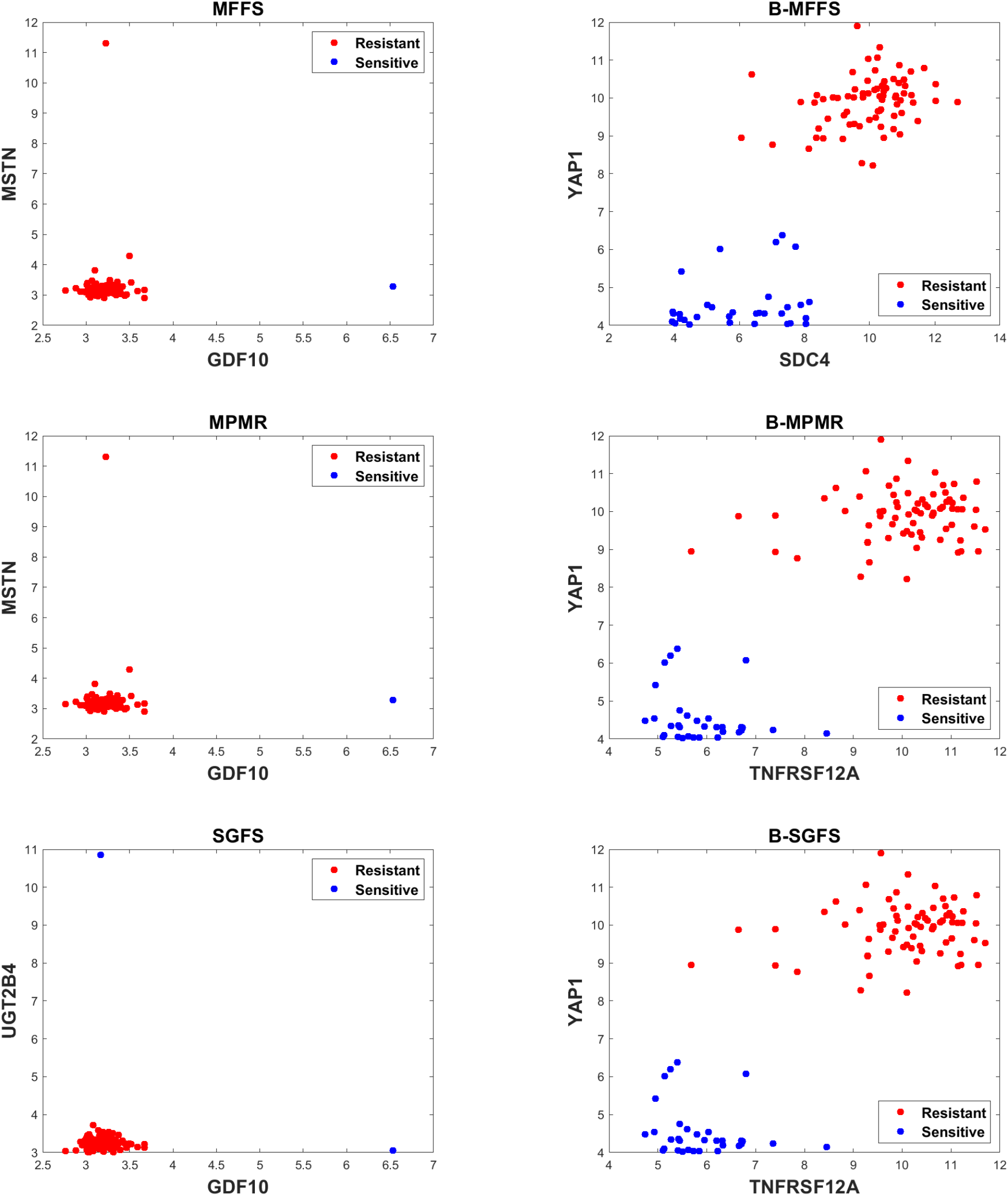
2-gene signatures selected by different feature selection methods.

## 6 Conclusion

High-dimensional data analysis is a foundation for systems approaches in medicine [74]. The applicability of these approaches in the real-world settings requires high dimensionality reduction [74]. This reduction helps to build predictive models in systems pharmacology of cancer to optimize therapeutic regimens proposed by *in vivo* and *in vitro* experiments [75]. Our results showed that our modified version of feature selection methods based on matrix factorization, we call B-MFFS, B-MPMR, and B-SGFS, could reduce the dimensionality of different datasets and provide feature signatures with higher accuracy than conventional methods. This is very important in the practice of quantitative and systems pharmacology, where we need to build both mechanistic and phenomenological models that emerge from omic data [76, 77]. To translate big complex data of omics and multi-omics into applied models for *in vitro, in vivo*, and clinical validation, we need such methodologies. This high dimensionality reduction can pave the path to establishing integrated systems pharmacology platforms that can predict drug responses in cancer cell lines and tumor models using small sets of bio-molecular markers [76].

One of the most potent applications of systems pharmacology is using dynamical models of signaling networks. These signaling networks consist of complex interactions among a large number of proteins or genes [78]. One of the primary practices to build and tune these models is to use prior biological knowledge to construct a reduced version of these interactions [78]. Our proposed methods can reduce these feature spaces quantitatively and select the most significant nodes in the interaction networks for more accurate and realistic dynamical models. The huge parameter spaces of these models also need reduction to represent a robust, optimized model [9]. Our dimensionality reduction methods can be applied to solve this problem as well.

Besides dynamic models, finding molecular sets to predict therapeutic responses in cancer models is an unmet need, and quantitative modalities like our proposed methodologies can provide the principles for this complex challenge. To improve the theoretical foundation of the proposed frameworks, it is important to determine that within a category of appropriate bases, which one is the most well-suited basis of features for a given unsupervised feature selection problem. This issue can be considered as a potential direction for future research.

## Author Contributions

A.M., I.T., F.S.M. and M.E., conceived the project. A.M., F.S.M., N.A., M.R.R., I.T., and M.E. designed the computational methodology. A.M., F.S.M., M.R.R., and I.T. did the computational analysis. All authors read and commented on the manuscript.

## Acknowledgments

A.M. and M.E. were supported by Iran National Science Foundation (INSF) and Shahid Bahonar University of Kerman [grant number: 97021946]. I.T. contributed to this paper at the Cellular Energetics Program of the Kavli Institute for Theoretical Physics, supported in part by the National Science Foundation Grant NSF PHY-1748958, NIH Grant R25GM067110, and the Gordon and Betty Moore Foundation Grant 2919.02.

## Abbreviations

ACC: Clustering Accuracy
CCLE: Cancer Cell Line Encyclopedia
DLBCL: Diffuse large B-cell lymphoma
GB: Glioblastoma Multiforme
LS: Laplacian Score
MBC: Molecular Biology of The Cell
MFFS: matrix factorization
MPMR: Maximum Projection and Minimum Redundancy
NMF: Nonnegative Matrix Factorization
NMI: Normalized Mutual Information
PCA: Principal Components Analysis
SGFS: Subspace learning-based Graph Regularized Feature Selection
SVD: Singular Value Decomposition
TKI: Tyrosine Kinase Inhibitor
TPR: True Positive Rate
TNR: True Negative Rate
VS: Variance Score

## Appendix

### A Global Conjugate Gradient Method

The global conjugate gradient method was developed as an extension of the conjugate gradient technique [54, 79]. Over the past recent years, it has been demonstrated that comparing to direct techniques such as the Gaussian elimination method, the iterative technique of global conjugate gradient enjoys a far better performance to deal with large matrix equations [80]. In such cases, it is usually supposed that constraints such as being both sparse and symmetric positive definite are imposed on the coefficient matrix corresponding to the given matrix equation [52]. A basic form of matrix equations whose exact solution can be estimated by this method is

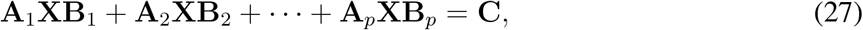

in which the matrices **A**_*j*_ ∈ ℝ^*n*×*n*^ and **B**_*j*_ ∈ ℝ^*m*×*m*^ are symmetric positive definite (or symmetric semi-positive definite) for *j* = 1, …, *p* and **C** ∈ ℝ^*n*×*m*^. The unknown matrix is also denoted by **X** ∈ ℝ^*n*×*m*^.

Algorithm A1 presents the global conjugate gradient framework.

#### Algorithm A1 The global conjugate gradient method.

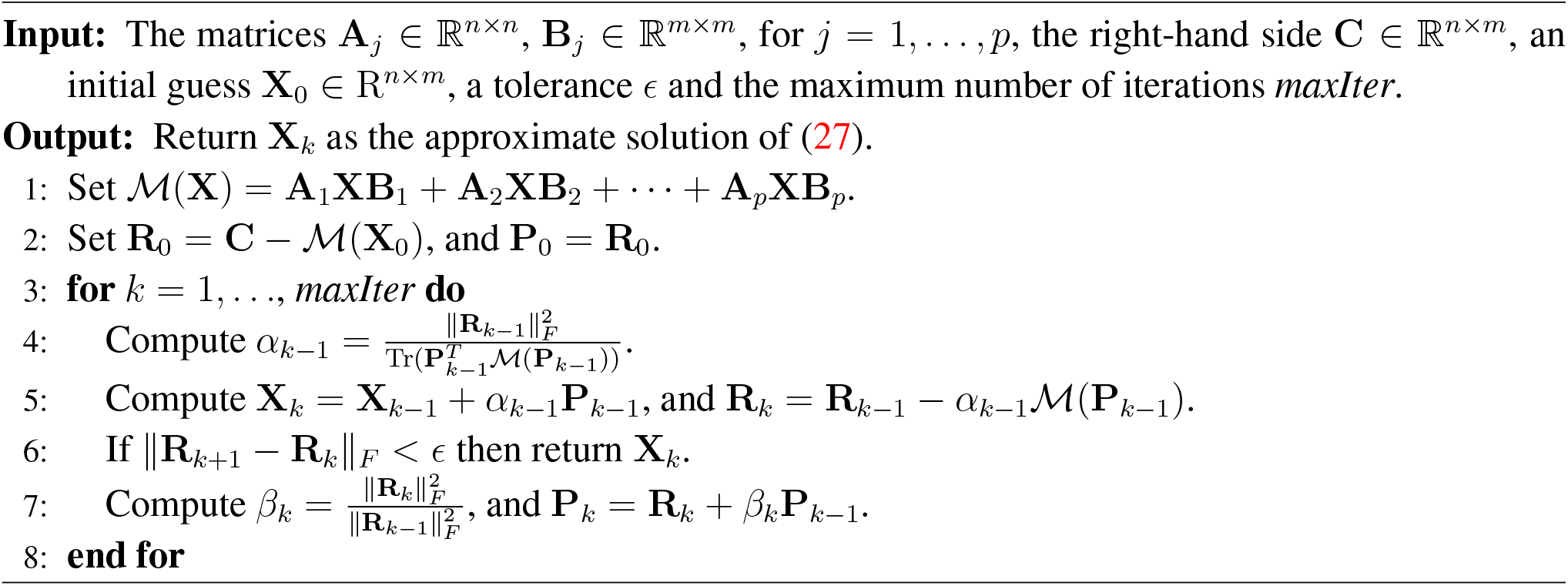

## References

[1] Iman Tavassoly, Joseph Goldfarb, and Ravi Iyengar. Systems biology primer: the basic methods and approaches. Essays in biochemistry, 62(4):487–500, 2018.

[2] Iman Tavassoly, Yuan Hu, Shan Zhao, Chiara Mariottini, Aislyn Boran, Yibang Chen, Lisa Li, Rosa E Tolentino, Gomathi Jayaraman, Joseph Goldfarb, et al. Genomic signatures defining responsiveness to allopurinol and combination therapy for lung cancer identified by systems therapeutics analyses. Molecular oncology, 13(8):1725–1743, 2019.

[3] Prashant Dogra, Joseph D Butner, Yao-li Chuang, Sergio Caserta, Shreya Goel, C Jeffrey Brinker, Vittorio Cristini, and Zhihui Wang. Mathematical modeling in cancer nanomedicine: a review. Biomedical Microdevices, 21(2):1–23, 2019.

[4] Shigeyuki Magi, Kazunari Iwamoto, and Mariko Okada-Hatakeyama. Current status of mathematical modeling of cancer–from the viewpoint of cancer hallmarks. Current Opinion in Systems Biology, 2:39–48, 2017.

[5] Miguel Ángel Medina. Mathematical modeling of cancer metabolism. Critical reviews in oncology/hematology, 124:37–40, 2018.

[6] Tayebeh Waezizadeh, Adel Mehrpooya, Maryam Rezaeizadeh, and Shantia Yarahmadian. Mathematical models for the effects of hypertension and stress on kidney and their uncertainty. Mathematical biosciences, 305:77–95, 2018.

[7] Iman Tavassoly. Dynamics of Cell Fate Decision Mediated by the Interplay of Autophagy and Apoptosis in Cancer Cells: Mathematical Modeling and Experimental Observations. Springer, 2015.

[8] Mohammadreza Dorvash, Mohammad Farahmandnia, and Iman Tavassoly. A systems biology roadmap to decode mTOR control system in cancer. Interdisciplinary Sciences: Computational Life Sciences, 12(1):1–11, 2020.

[9] I Tavassoly, J Parmar, AN Shajahan-Haq, R Clarke, William T Baumann, and John J Tyson. Dynamic modeling of the interaction between autophagy and apoptosis in mammalian cells. CPT: pharmacometrics & systems pharmacology, 4(4):263–272, 2015.

[10] Evanthia Koukouli, Dennis Wang, Frank Dondelinger, and Juhyun Park. A regularized functional regression model enabling transcriptome-wide dosage-dependent association study of cancer drug response. PLoS computational biology, 17(1):e1008066, 2021.

[11] Mehrdad Rostami, Kamal Berahmand, Elahe Nasiri, and Saman Forouzande. Review of swarm intelligence-based feature selection methods. Engineering Applications of Artificial Intelligence, 100:104210, 2021.

[12] Dong Wang, Jin-Xing Liu, Ying-Lian Gao, Chun-Hou Zheng, and Yong Xu. Characteristic gene selection based on robust graph regularized non-negative matrix factorization. IEEE/ACM transactions on computational biology and bioinformatics, 13(6):1059–1067, 2015.

[13] Jean-Philippe Brunet, Pablo Tamayo, Todd R Golub, and Jill P Mesirov. Metagenes and molecular pattern discovery using matrix factorization. Proceedings of the national academy of sciences, 101(12):4164–4169, 2004.

[14] Karthik Devarajan. Nonnegative matrix factorization: an analytical and interpretive tool in computational biology. PLoS Comput Biol, 4(7):e1000029, 2008.

[15] Raffaele Giancarlo and Filippo Utro. Speeding up the consensus clustering methodology for microarray data analysis. Algorithms for Molecular Biology, 6(1):1–13, 2011.

[16] Belhassen Bayar, Nidhal Bouaynaya, and Roman Shterenberg. Probabilistic non-negative matrix factorization: theory and application to microarray data analysis. Journal of bioinformatics and computational biology, 12(01):1450001, 2014.

[17] Isabelle Guyon, Jason Weston, Stephen Barnhill, and Vladimir Vapnik. Gene selection for cancer classification using support vector machines. Machine learning, 46(1):389–422, 2002.

[18] Yue Li and Zhaolei Zhang. Computational biology in microRNA. Wiley Interdisciplinary Reviews: RNA, 6(4):435–452, 2015.

[19] Jin-Xing Liu, Yong Xu, Chun-Hou Zheng, Heng Kong, and Zhi-Hui Lai. Rpca-based tumor classification using gene expression data. IEEE/ACM Transactions on Computational Biology and Bioinformatics, 12(4):964–970, 2014.

[20] Haifa Ben Saber and Mourad Elloumi. DNA microarray data analysis: a new survey on biclustering. International Journal for Computational Biology (IJCB), 4(1):21–37, 2015.

[21] Chun-Hou Zheng, Lei Zhang, To-Yee Ng, Chi Keung Shiu, and De-Shuang Huang. Metasample-based sparse representation for tumor classification. IEEE/ACM Transactions on Computational Biology and Bioinformatics, 8(5):1273–1282, 2011.

[22] Verónica Bolón-Canedo, Noelia Sánchez-Maroño, and Amparo Alonso-Betanzos. A review of feature selection methods on synthetic data. Knowledge and information systems, 34(3):483–519, 2013.

[23] Isabelle Guyon, Steve Gunn, Masoud Nikravesh, and Lofti A Zadeh. Feature extraction: foundations and applications, volume 207. Springer, 2008.

[24] Zena M Hira and Duncan F Gillies. A review of feature selection and feature extraction methods applied on microarray data. Advances in bioinformatics, 2015, 2015.

[25] Alexandros Kalousis, Julien Prados, and Melanie Hilario. Stability of feature selection algorithms: a study on high-dimensional spaces. Knowledge and information systems, 12(1):95–116, 2007.

[26] Mehrdad Rostami, Saman Forouzandeh, Kamal Berahmand, and Mina Soltani. Integration of multi-objective PSO based feature selection and node centrality for medical datasets. Genomics, 112(6):4370–4384, 2020.

[27] Jun Chin Ang, Andri Mirzal, Habibollah Haron, and Haza Nuzly Abdull Hamed. Supervised, unsupervised, and semi-supervised feature selection: a review on gene selection. IEEE/ACM transactions on computational biology and bioinformatics, 13(5):971–989, 2015.

[28] Verónica Bolón-Canedo, Noelia Sánchez-Marono, Amparo Alonso-Betanzos, José Manuel Benítez, and Francisco Herrera. A review of microarray datasets and applied feature selection methods. Information Sciences, 282:111–135, 2014.

[29] Mahla Mokhtia, Mahdi Eftekhari, and Farid Saberi-Movahed. Feature selection based on regularization of sparsity based regression models by hesitant fuzzy correlation. Applied Soft Computing, 91:106255, 2020.

[30] Jie Cai, Jiawei Luo, Shulin Wang, and Sheng Yang. Feature selection in machine learning: A new perspective. Neurocomputing, 300:70–79, 2018.

[31] Mahdi Eftekhari, Farid Saberi-Movahed, and Adel Mehrpooya. Supervised feature selection via information gain, maximum projection and minimum redundancy. In SLAA10 Seminar Linear Algebra and Its Application, pages 29–35, 2020.

[32] Ronghua Shang, Jiuzheng Song, Licheng Jiao, and Yangyang Li. Double feature selection algorithm based on low-rank sparse non-negative matrix factorization. International Journal of Machine Learning and Cybernetics, pages 1–18, 2020.

[33] Jianglin Lu, Hailing Wang, Jie Zhou, Yudong Chen, Zhihui Lai, and Qinghua Hu. Low-rank adaptive graph embedding for unsupervised feature extraction. Pattern Recognition, 113:107758, 2021.

[34] Miao Qi, Ting Wang, Fucong Liu, Baoxue Zhang, Jianzhong Wang, and Yugen Yi. Unsupervised feature selection by regularized matrix factorization. Neurocomputing, 273:593–610, 2018.

[35] Farid Saberi-Movahed, Mahdi Eftekhari, and Mohammad Mohtashami. Supervised feature selection by constituting a basis for the original space of features and matrix factorization. International Journal of Machine Learning and Cybernetics, 11:1405–1421, 2020.

[36] Ronghua Shang, Kaiming Xu, Fanhua Shang, and Licheng Jiao. Sparse and low-redundant subspace learning-based dual-graph regularized robust feature selection. Knowledge-Based Systems, 187:104830, 2020.

[37] Shiping Wang, Witold Pedrycz, Qingxin Zhu, and William Zhu. Subspace learning for unsupervised feature selection via matrix factorization. Pattern Recognition, 48(1):10–19, 2015.

[38] Shiping Wang, Witold Pedrycz, Qingxin Zhu, and William Zhu. Unsupervised feature selection via maximum projection and minimum redundancy. Knowledge-Based Systems, 75:19–29, 2015.

[39] Ronghua Shang, Wenbing Wang, Rustam Stolkin, and Licheng Jiao. Subspace learning-based graph regularized feature selection. Knowledge-Based Systems, 112:152–165, 2016.

[40] Saúl Solorio-Fernández, J Ariel Carrasco-Ochoa, and José Fco Martínez-Trinidad. A review of unsupervised feature selection methods. Artificial Intelligence Review, 53(2):907–948, 2020.

[41] Emrah Hancer, Bing Xue, and Mengjie Zhang. A survey on feature selection approaches for clustering. Artificial Intelligence Review, 53(6):4519–4545, 2020.

[42] Benjamin Auffarth, Maite López, and Jesús Cerquides. Comparison of redundancy and relevance measures for feature selection in tissue classification of ct images. In Industrial conference on data mining, pages 248–262. Springer, 2010.

[43] Charu C Aggarwal. Linear Algebra and Optimization for Machine Learning. Springer, 2020.

[44] Carl D Meyer. Matrix analysis and applied linear algebra, volume 71. SIAM, 2000.

[45] Gene H Golub and Christian Reinsch. Singular value decomposition and least squares solutions. In Linear algebra, pages 134–151. Springer, 1971.

[46] Daniel D Lee and H Sebastian Seung. Learning the parts of objects by non-negative matrix factorization. Nature, 401(6755):788–791, 1999.

[47] I.T. Jolliffe. Principal Component Analysis. Springer, 1986.

[48] Feiping Nie, Heng Huang, Xiao Cai, and Chris Ding. Efficient and robust feature selection via joint l_2,1_-norm minimization. Advances in neural information processing systems, 23:1813–1821, 2010.

[49] Khadidja Henni, Neila Mezghani, and Charles Gouin-Vallerand. Unsupervised graph-based feature selection via subspace and pagerank centrality. Expert Systems with Applications, 114:46–53, 2018.

[50] Shaoyong Li, Chang Tang, Xinwang Liu, Yaping Liu, and Jiajia Chen. Dual graph regularized compact feature representation for unsupervised feature selection. Neurocomputing, 331:77–96, 2019.

[51] Chang Tang, Meiru Bian, Xinwang Liu, Miaomiao Li, Hua Zhou, Pichao Wang, and Hailin Yin. Unsupervised feature selection via latent representation learning and manifold regularization. Neural Networks, 117:163–178, 2019.

[52] André Gaul. Recycling Krylov subspace methods for sequences of linear systems: Analysis and applications. PhD thesis, Technischen Universitat Berlin, 2014.

[53] A El Guennouni, Khalide Jbilou, and AJ Riquet. Block Krylov subspace methods for solving large Sylvester equations. Numerical Algorithms, 29(1-3):75–96, 2002.

[54] Mohammed Heyouni and Azeddine Essai. Matrix Krylov subspace methods for linear systems with multiple right-hand sides. Numerical Algorithms, 40(2):137–156, 2005.

[55] Scott L Pomeroy, Pablo Tamayo, Michelle Gaasenbeek, Lisa M Sturla, Michael Angelo, Margaret E McLaughlin, John YH Kim, Liliana C Goumnerova, Peter M Black, Ching Lau, et al. Prediction of central nervous system embryonal tumour outcome based on gene expression. Nature, 415(6870):436–442, 2002.

[56] Uri Alon, Naama Barkai, Daniel A Notterman, Kurt Gish, Suzanne Ybarra, Daniel Mack, and Arnold J Levine. Broad patterns of gene expression revealed by clustering analysis of tumor and normal colon tissues probed by oligonucleotide arrays. Proceedings of the National Academy of Sciences, 96(12):6745–6750, 1999.

[57] Ash A Alizadeh, Michael B Eisen, R Eric Davis, Chi Ma, Izidore S Lossos, Andreas Rosenwald, Jennifer C Boldrick, Hajeer Sabet, Truc Tran, Xin Yu, et al. Distinct types of diffuse large B-cell lymphoma identified by gene expression profiling. Nature, 403(6769):503–511, 2000.

[58] William A Freije, F Edmundo Castro-Vargas, Zixing Fang, Steve Horvath, Timothy Cloughesy, Linda M Liau, Paul S Mischel, and Stanley F Nelson. Gene expression profiling of gliomas strongly predicts survival. Cancer research, 64(18):6503–6510, 2004.

[59] Jundong Li, Kewei Cheng, Suhang Wang, Fred Morstatter, Robert P Trevino, Jiliang Tang, and Huan Liu. Feature selection: A data perspective. ACM Computing Surveys (CSUR), 50(6):94, 2018.

[60] Jordi Barretina, Giordano Caponigro, Nicolas Stransky, Kavitha Venkatesan, Adam A Margolin, Sungjoon Kim, Christopher J Wilson, Joseph Lehár, Gregory V Kryukov, Dmitriy Sonkin, et al. The cancer cell line encyclopedia enables predictive modelling of anticancer drug sensitivity. Nature, 483(7391):603–607, 2012.

[61] Christopher M Bishop et al. Neural networks for pattern recognition. Oxford university press, 1995.

[62] Xiaofei He, Deng Cai, and Partha Niyogi. Laplacian score for feature selection. Advances in neural information processing systems, 18:507–514, 2005.

[63] Verónica Bolón-Canedo, Noelia Sánchez-Maroño, and Amparo Alonso-Betanzos. On the effectiveness of discretization on gene selection of microarray data. In The 2010 International Joint Conference on Neural Networks (IJCNN), pages 1–8. IEEE, 2010.

[64] Yvan Saeys, Inaki Inza, and Pedro Larranaga. A review of feature selection techniques in bioinformatics. bioinformatics, 23(19):2507–2517, 2007.

[65] Mohammad Najafzadeh, Amir Etemad-Shahidi, and Siow Yong Lim. Scour prediction in long contractions using ANFIS and SVM. Ocean Engineering, 111:128–135, 2016.

[66] Katherine M Aird, Jennifer L Allensworth, Ines Batinic-Haberle, H Kim Lyerly, Mark W De-whirst, and Gayathri R Devi. ErbB1/2 tyrosine kinase inhibitor mediates oxidative stress-induced apoptosis in inflammatory breast cancer cells. Breast cancer research and treatment, 132(1):109–119, 2012.

[67] Marija Mihajlovic, Branka Ivkovic, Biljana Jancic-Stojanovic, Aleksandra Zeljkovic, Vesna Spasojevic-Kalimanovska, Jelena Kotur-Stevuljevic, and Dragana Vujanovic. Modulation of oxidative stress/antioxidative defence in human serum treated by four different tyrosine kinase inhibitors. Anti-cancer drugs, 31(9):942–949, 2020.

[68] Mohamed E Shaker. Nilotinib interferes with the signalling pathways implicated in acetaminophen hepatotoxicity. Basic & clinical pharmacology & toxicology, 114(3):263–270, 2014.

[69] Jens Hansen, David Meretzky, Simeneh Woldesenbet, Gustavo Stolovitzky, and Ravi Iyengar. A flexible ontology for inference of emergent whole cell function from relationships between subcellular processes. Scientific reports, 7(1):1–13, 2017.

[70] Jesper B Andersen, Bart Spee, Boris R Blechacz, Itzhak Avital, Mina Komuta, Andrew Barbour, Elizabeth A Conner, Matthew C Gillen, Tania Roskams, Lewis R Roberts, et al. Genomic and genetic characterization of cholangiocarcinoma identifies therapeutic targets for tyrosine kinase inhibitors. Gastroenterology, 142(4):1021–1031, 2012.

[71] Soon Auck Hong, Si-Hyong Jang, Mee-Hye Oh, Sung Joon Kim, Jin-Hyung Kang, and Sook-Hee Hong. Overexpression of YAP1 in EGFR mutant lung adenocarcinoma prior to tyrosine kinase inhibitor therapy is associated with poor survival. Pathology-Research and Practice, 214(3):335–342, 2018.

[72] Jai-Nien Tung, Po-Lin Lin, Yao-Chen Wang, De-Wei Wu, Chi-Yi Chen, and Huei Lee. Pd-l1 confers resistance to EGFR mutation-independent tyrosine kinase inhibitors in non-small cell lung cancer via upregulation of YAP1 expression. Oncotarget, 9(4):4637, 2018.

[73] Timothy G Whitsett, Emily Cheng, Landon Inge, Kaushal Asrani, Nathan M Jameson, Galen Hostetter, Glen J Weiss, Christopher B Kingsley, Joseph C Loftus, Ross Bremner, et al. Elevated expression of Fn14 in non-small cell lung cancer correlates with activated EGFR and promotes tumor cell migration and invasion. The American journal of pathology, 181(1):111–120, 2012.

[74] Charles Auffray, Zhu Chen, and Leroy Hood. Systems medicine: the future of medical genomics and healthcare. Genome medicine, 1(1):1–11, 2009.

[75] Iman Tavassoly, Yuan Hu, Shan Zhao, Chiara Mariottini, Aislyn Boran, Yibang Chen, Lisa Li, Rosa E Tolentino, Gomathi Jayaraman, Joseph Goldfarb, et al. Genomic signatures defining responsiveness to allopurinol and combination therapy for lung cancer identified by systems therapeutics analyses. Molecular oncology, 13(8):1725–1743, 2019.

[76] Iman Tavassoly, Joseph Goldfarb, and Ravi Iyengar. Systems biology primer: the basic methods and approaches. Essays in biochemistry, 62(4):487–500, 2018.

[77] Piet H Van der Graaf and Neil Benson. Systems pharmacology: bridging systems biology and pharmacokinetics-pharmacodynamics (PKPD) in drug discovery and development. Pharmaceutical research, 28(7):1460–1464, 2011.

[78] John J Tyson, William T Baumann, Chun Chen, Anael Verdugo, Iman Tavassoly, Yue Wang, Louis M Weiner, and Robert Clarke. Dynamic modelling of oestrogen signalling and cell fate in breast cancer cells. Nature Reviews Cancer, 11(7):523–532, 2011.

[79] Valeria Simoncini. Computational methods for linear matrix equations. SIAM Review, 58(3):377–441, 2016.

[80] Mohammed Heyouni, Farid Saberi-Movahed, and Azita Tajaddini. On global Hessenberg based methods for solving Sylvester matrix equations. Computers & Mathematics with Applications, 77(1):77–92, 2019.

